# Thermal and mechanical modalities converge in the noxious range

**DOI:** 10.1101/2025.03.03.641313

**Authors:** Feng Wang, Samuel Ferland, Sylvain L. Côté, Louis-Étienne Lorenzo, Erik Bélanger, Pauline Larqué, Changlin Li, Antoine Godin, Bo Duan, Daniel C. Côté, Marie-Eve Paquet, Yves De Koninck

## Abstract

There are competing theories on how nociceptive input is encoded by sensory fibers. That sensory modalities are exclusively conveyed by distinct populations of fibers is supported by ablation studies, indicating that TRPV1^+^ and MrgprD^+^ afferents selectively encode for heat and mechanical input, respectively. However, interpreting ablation results can be clouded by compensatory plasticity. Furthermore, a population-level parametric analysis of afferent response profiles to natural stimuli *in vivo* is still scarce. Using functional imaging *in vivo* in mice, we found that most TRPV1^+^ neurons responded to noxious heat, but not cooling. While none of them responded to innocuous mechanical input, about half were also sensitive to noxious mechanical stimuli. In contrast, 80% of MrgprD^+^ neurons responded to noxious mechanical stimuli. Yet, 70% were also sensitive to noxious thermal stimuli. Of innocuous mechanosensitive MrgprD^+^ neurons, <15% responded to innocuous warming and none to innocuous cooling. Polymodality in the innocuous thermal and mechanical range also occurred in <10% of all primary afferents. Acute silencing of TRPV1^+^ or MrgprD^+^ afferents inhibited both thermal and mechanical nociception. In contrast, ablation did not reproduce a combined loss of sensitivity, which appeared to be due to compensatory central disinhibition. Our findings reveal that modality separation dominates in the innocuous regime, while polymodality predominates in the noxious range.

## Introduction

Pain is “an unpleasant sensory and emotional experience associated with actual or potential tissue damage” ^1^. Many types of pain have been classified using different criteria. For example, pain can be separated into nociceptive pain, inflammatory pain, and neuropathic pain based on whether the pain is protective or maladaptive ^2–4^. In human studies, pathological pain categories tend to be based on the perceived location (e.g. low back pain and migraine) or etiology (e.g. cancer pain and neuropathic pain) ^5^, rather than modality. In contrast, in animal studies, withdrawal reflex from presumed noxious stimuli is often used as a proxy for pain, and the type of pain is classified as mechanical, thermal, or chemical pain according to the modality of stimulus eliciting the reflex ^6,7^. These two angles of classification may converge, as in some studies with large multi-aetiology clinical cohorts, regardless of their underlying disease, most neuropathic pain patients were classified into one of three distinct sensory subgroups: those with sensory loss, mechanical hyperalgesia, or thermal hyperalgesia ^8^. Yet it remains uncertain if this is related to the innocuous cue associated with the stimuli or whether the patients can truly distinguish the painful modality.

The concept that pain can be categorized on the basis of modalities is, in fact, mainly an extrapolation from animal studies ^6,9,10^. Nociceptors in mice are almost purely separated into peptidergic *vs.* non-peptidergic primary afferents, based on whether they express neuropeptides, such as CGRP, or not ^11,12^. Ablating most TRPV1^+^ afferents, a large subpopulation of peptidergic neurons, with intrathecal injection of high-dose capsaicin or resiniferatoxin (specific agonists of TRPV1) led to a complete loss of heat sensitivity, while the mechanical sensitivity was not affected ^13–15^, suggesting that TRPV1^+^ dorsal root ganglion (DRG) neurons are essential for thermal nociception, but not mechanical nociception. In contrast, ablation of MrgprD^+^ neurons, a large subpopulation of non-peptidergic DRG neurons, only attenuated withdrawal to noxious mechanical stimuli, but not to thermal stimuli ^13^. These findings appear to support a labeled line framework for sensory coding ^16–18^. Yet, they still do not allow us to determine whether the animals can *distinguish* nociceptive modalities; let alone distinguish pain sensation on the basis of modalities.

Contrasting findings emerge from physiological studies of DRG neurons, arguing against the concept of labeled lines ^17–19^. *Ex-vivo* recording of DRG neurons combined with labeling showed that TRPV1^+^ neurons only responded to noxious heat stimulation, but not to noxious pinching stimulation ^20,21^. However, IB4^+^ non-peptidergic neurons were sensitive to both noxious thermal and mechanical stimulation ^20^. Moreover, a substantial percentage of nociceptors were found to be polymodal in both electrophysiological ^22,23^ and *in vivo* imaging studies ^24,25^. These findings tend to favor a combinatorial framework for sensory coding ^17–19,26,27^.

In the current study, we sought to reconcile the discrepancy between physiological and behavioral ablation studies. To achieve this, we conducted functional characterization of genetically identified populations of TRPV1^+^ and MrgprD^+^ neurons *in vivo* combined with behavioral assessment after respectively blocking their activity acutely. Our results revealed that about half of TRPV1^+^ and MrgprD^+^ neurons overlap in terms of modalities in the noxious regime, but not in the innocuous regime. Furthermore, silencing either class of afferent, individually, attenuates in each case nocifensive response to both noxious thermal and mechanical stimuli, suggesting that while modality separation appears to dominate in the innocuous regime, thermal and mechanical modalities merge in the noxious range.

## Results

### TRPV1^+^ sensory neurons encode noxious heat in a graded fashion

We sought to address two questions regarding TRPV1^+^ neurons: first, whether under *in vivo* conditions, they are noxious heat specific and, second, whether they form the only population sensitive to noxious heat. To address the first question, we performed *in vivo* Ca^+^ imaging from TRPV1^+^ DRG neurons. We injected an adeno-associated virus (AAV) carrying Cre-dependent GCaMP6s ^28^ into newborn TRPV1^Cre^ knockin mice ^29^. Injection in heterozygous TRPV1^Cre/+^ mice resulted in robust expression of GCaMP in about half of TRPV1^+^ DRG neurons (Fig. 1a, b). The transduction efficiency was similar to that previously reported ^25^. The viral transduction only labeled Cre^+^ neurons, as no GCaMP labeled neuron was found in wild-type mice (Supplementary Fig. 1).

**Figure 1.**
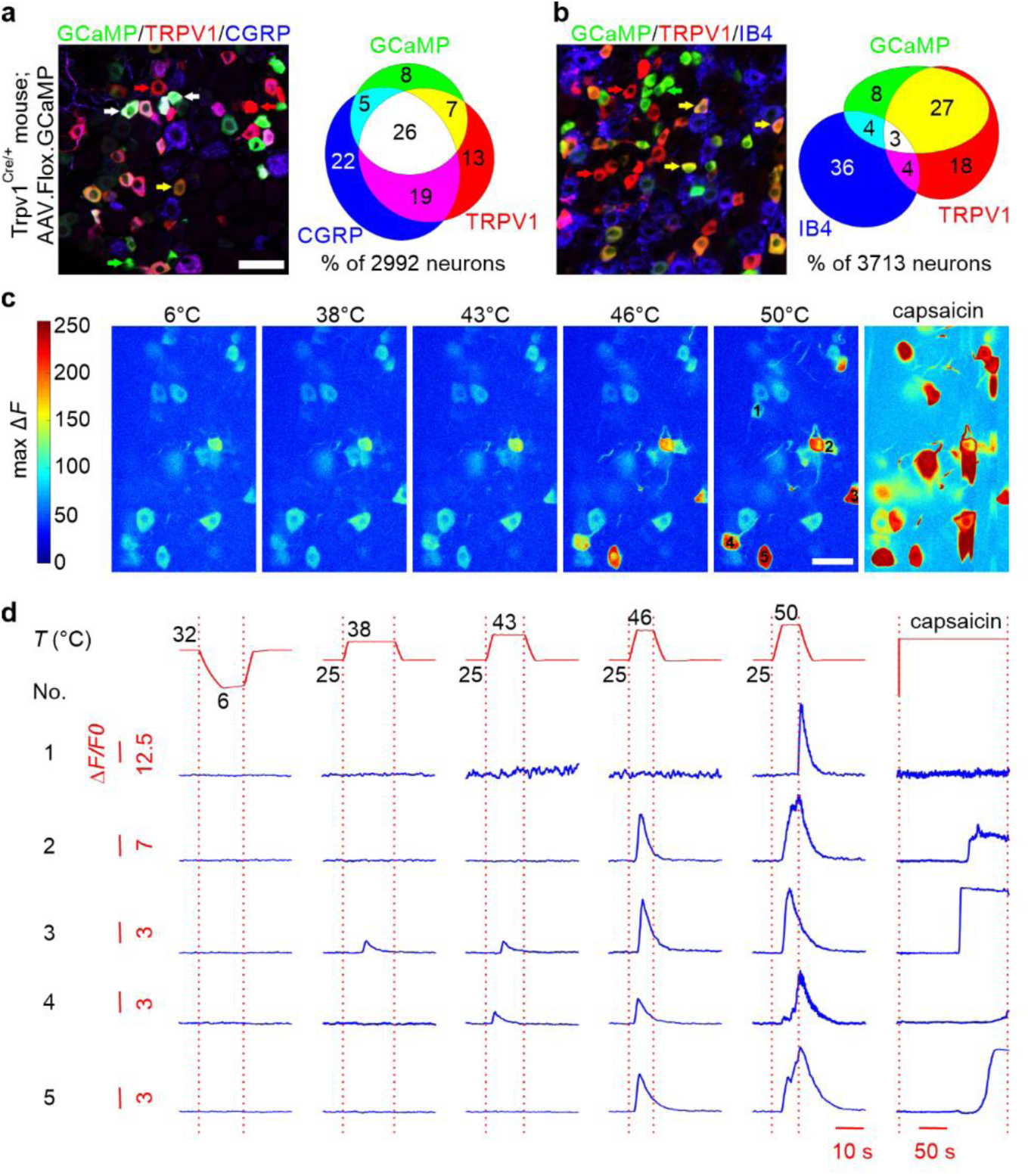
*In vivo* functional imaging of TRPV1^+^ sensory afferents. (a-b) Representative micrographs of DRG sections from Trpv1^Cre/+^ mice injected with AAV showing GCaMP6s expression (without antibody amplification) superimposed on labeling for TRPV1 and peptidergic neuron marker, CGRP (a), as well as TRPV1 and non-peptidergic neuron marker, IB4 (b). The white arrows pointed to the neurons that were triple-positive, the yellow arrows pointed to the neurons that were GCaMP6s and TRPV1-double positive, while the green and red arrows pointed to the neurons that were only GCaMP6s or TRPV1-positive, respectively. Euler diagrams showed that the majority of GCaMP^+^ neurons expressed TRPV1. Scale bar = 50 µm. (c) Color-coded heatmaps of a typical imaging field showing neuronal responses to noxious and innocuous thermal stimuli in Trpv1^Cre/+^; AAV.Flex.GCaMP mouse. Color scale indicates maximum △*F*. Scale bar = 50 µm. (d) Representative Ca^2+^ curves from five neurons, highlighted in C, showing typical responses to thermal and capsaicin stimulation. All five neurons, except neuron 1, were sensitive to capsaicin stimulation. Note the different Δ*F*/*F0* scales for different neurons. The upper panels show the protocols for fast-ramp-and-hold thermal stimuli applied to the plantar side of hind paw, as well as the bath application of capsaicin. A baseline of 32°C and 25°C was chosen for cooling and heating stimuli, respectively.

TRPV1 expression is reduced and refined during the development ^29^. In adult wild-type mice, we found that the majority of TRPV1^+^ neurons are CGRP^+^ peptidergeric neurons, but not IB4^+^ non-peptidergic neurons (Supplementary Fig. 2), consistent with previous reports ^20,29^. In AAV transduced heterozygous TRPV1^Cre/+^ mice, > 70% of GCaMP^+^ neurons were TRPV1^+^, but not IB4^+^ (Fig. 1a, b), proving that most of GCaMP labeled neurons expressed TRPV1. On the other hand, in homozygous TRPV1^Cre/Cre^ mice, only about 50% of GCaMP labeled neurons expressed TRPV1, and about 30% of GCaMP^+^ neurons were also IB4^+^ (Supplementary Fig. 3). Thus, only heterozygous TRPV1^Cre/+^ mice were used for the imaging experiment. Furthermore, only neurons with pharmacologically confirmed TRPV1 expression *post-hoc* were included in the analysis (see below).

Then, we performed *in vivo* Ca^2+^ imaging from L4 DRG on anesthetized mice using two-photon microscopy ^25^ and applied a series of thermal stimuli to the plantar area of the hind paw, ranging from noxious cold to noxious heat (Fig. 1c, d). Robust Ca^2+^ responses were observed when noxious heat (50°C) was applied. In some neurons, warm (38°C) and heat (43°C) also induced a smaller Ca^2+^ response (Fig. 1c, d). Towards the end of the imaging session, we also applied capsaicin directly to the exposed DRG to validate the presence of TRPV1 channel; most GCaMP^+^ thermoreceptors responded to capsaicin (Fig. 1c, d). As mentioned above, to exclude GCaMP^+^, but TRPV1 negative neurons, we only included capsaicin responsive neurons for further analysis.

Among the one thousand neurons imaged, 125 neurons showed clear thermal responses delivered to the hind paw, and 99 (80%) of these were confirmed capsaicin-sensitive and included in the analysis. These TRPV1^+^ neurons did not respond to noxious cold (6°C), cold (15°C), or innocuous cool (20°C) stimuli (Fig. 2a). While 20% of neurons responded to innocuous warm (38°C), a greater percentage responded to noxious heat (43-46°C), and almost all the neurons (95%) responded to 50°C (Fig. 2a).

**Figure 2.**
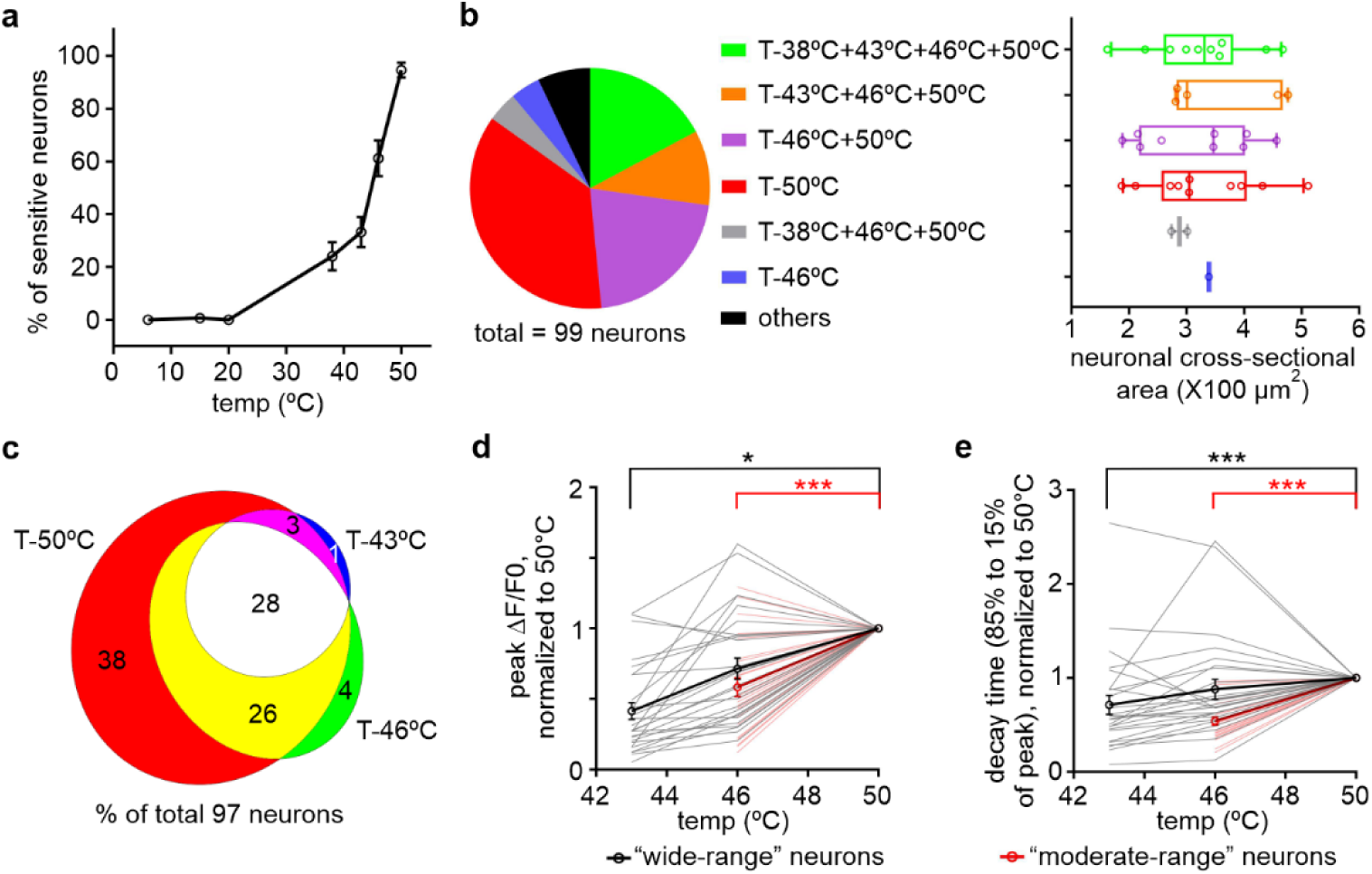
TRPV1^+^ sensory afferents encode heat temperature in a graded fashion at both population and single-neuron levels. (a) Proportion of thermo-sensitive TRPV1^+^ neurons (*i.e.* that responded to at least one type of thermal stimulus applied to the plantar side of hind paw) in each imaging field that responded to different thermal stimuli (n = 11 fields from 11 mice). (b) Left: pie chart showing the distribution of TRPV1^+^ neurons with distinct thermal response profiles. Right: distribution of neuronal size for each of the subgroups defined on the left (n = 1-10 neurons per subgroup). (c) Euler diagram showing overlap in neurons responsive to the heat/noxious heat stimuli. Numbers indicated the percentage of the neurons responding to at least one of the three stimuli as indicated. (d-e) Plots of normalized peak amplitude (d) and decay time (85% to 15% of peak; e) of “wide-range” heat-sensitive neurons (gray curves = individual neuron response; black curve = mean response of the group), as well as “moderate-range” noxious heat-sensitive neurons (pale red curves = individual neuron response; red curve = mean response of the group), to increasing temperature. Both “wide-range” heat and “moderate-range” noxious heat neurons displayed significantly greater and longer responses as temperature increased (n = 25-27 cells, *p < 0.05 and ***p < 0.001 using One-way Repeated Measures ANOVA test followed with Tukey’s multiple comparisons test for “wide-range” heat neurons and paired *t* test for “moderate-range” noxious heat neurons).

Then we categorized these neurons into different groups based on their thermal sensitivities. The largest group was composed of neurons responding only to 50°C; neurons responded to both 46°C and 50°C comprised the second largest group. The neurons responding to temperatures >43°C or 38°C also formed large groups. These four groups comprised around 85% of all TRPV1^+^ thermoreceptors. The remaining groups (each < 3%) were lumped together in ‘others’ (Fig. 2b).

The soma size of these neurons was also measured *in vivo*, and most of them were small-diameter neurons (Fig. 2b), consistent with histological findings (Fig. 1a) and previous reports that most TRPV1^+^ neurons are small-diameter peptidergic neurons ^29^. Also, we did not observe any clear difference in the soma size among neurons with different thermal sensitivity profiles (Fig. 2b; n = 1-10 cells, p = 0.95).

We analyzed the distribution of noxious heat-sensitive TRPV1^+^ neurons (> 43°C). We found that neurons responding to lower temperature formed nested subsets to the neurons responding to higher temperature (Fig. 2c). This indicated graded coding of intensity at the population level, as we previously reported ^25^.

To test whether individual TRPV1^+^ neurons encoded noxious heat intensity, we analysed the responses of neurons sensitive to a wide (43-50°C) *vs.* a moderate range (46-50°C) of temperatures. Both types displayed larger peak amplitude and longer decay at higher temperatures (Fig. 2d, e; n = 26-27 cells, *p < 0.05 and ***p < 0.001), indicating that individual TRPV1^+^ neurons encode noxious heat in a graded fashion as we previously reported ^25^.

Thus, while a significant subset of TRPV1^+^ thermoreceptors responds to innocuous warm, the majority are heat nociceptors, encoding heat in a graded fashion as individual neurons and as a population.

### Half of TRPV1^+^ neurons are polymodal nociceptors

To address the question of whether TRPV1^+^ neurons are heat-specific, we tested their mechanosensitivity in the innocuous and noxious ranges. Consistent with them being nociceptors, no TRPV1^+^ neurons responded to gentle brushing stimulation of the hind paw (Fig. 3a, b, d). Surprisingly, however, a large subpopulation of them (40%) displayed robust responses to noxious pinch (Fig. 3a, c, d), making them polymodal nociceptors. And while some pinch-sensitive neurons did not respond to 50°C stimulation of the plantar side, virtually all of them responded from either the plantar or dorsal side (Fig. 3e), confirming that all TRPV1^+^ neurons were heat nociceptors. Furthermore, the average peak amplitude of the pinch responses was even larger (133%) than that of 50°C responses (Fig. 3f), confirming that a large subpopulation of TRPV1^+^ neurons are polymodal nociceptors. Finally, we used single-cell PCR to examine whether TRPV1^+^ neurons express Piezo2, which appears essential for touch sensation and mechanonociception ^30–32^. Surprisingly, all the TRPV1^+^ neurons tested were Piezo2 positive (Fig. 3g), indicating that TRPV1^+^ neurons indeed have the machinery for mechanosensation.

**Figure 3.**
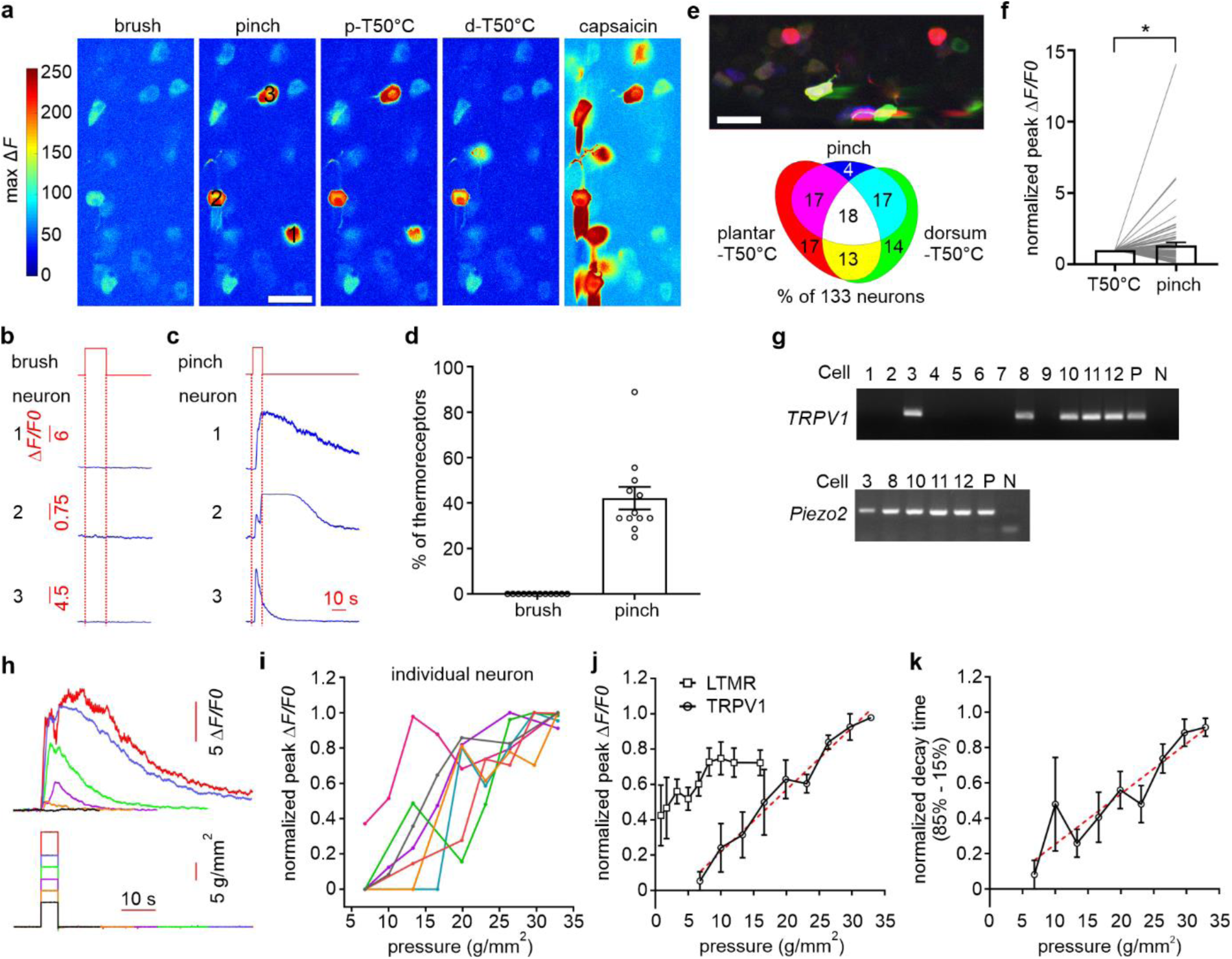
TRPV1^+^ sensory afferents are sensitive to noxious mechanical stimuli. (a) Color-coded heatmap of a typical imaging field showing neuronal responses to mechanical, thermal, and capsaicin stimulation in Trpv1^Cre/+^; AAV.Flex.GCaMP mouse. The color scale represents maximum △*F*. The noxious heat stimulation was applied to both plantar (p-T50°C) and dorsal side (d-T50°C) of hind paw in separate trials. Scale bar = 50 µm. (b-c) Representative Ca^2+^ curves from three neurons, highlighted in (A), show typical responses to pinching stimulation (C) and no response to brushing stimulation (B). Note the different Δ*F*/*F0* scales for different neurons. Neurons 2 and 3, but not neuron 1, were sensitive to capsaicin stimulation. The upper panels show the onset and offset of mechanical stimulation. Multiple repeated brushing or pinching stimulation was applied in each trial. (d) Proportion of thermo-sensitive TRPV1^+^ neurons in each imaging field tested with thermal stimulation on the plantar side of the hind paw that responded to brushing or pinching stimuli. = 11 fields from 11 mice. (e) Top: Color-coded heatmap of a typical imaging field showing neuronal responses to noxious heat stimuli applied to the dorsal (green) and plantar (red) sides of the hind paw, as well as to pinch stimuli (blue) applied to both sides. The intensity level of each channel represents the neuronal activation level to the corresponding stimulus. Scale bar = 50 µm. Bottom: Euler diagram showing overlap in neurons sensitive to noxious heat and pinch stimuli. Numbers indicated the percentage of the neurons responding to at least one of the three stimuli as indicated. (f) TRPV1^+^ neurons displayed slightly greater responses to pinching stimulation than noxious heat (n = 75 cells, *p < 0.05 using paired *t* test). (g) Single-cell PCR showed that most TRPV1^+^ DRG neurons express mechanoreceptor Piezo2. P is positive control and N is negative control. (h) Representative Ca^2+^ curves from one TRPV1^+^ DRG neuron to a series of indentation stimuli with increasing pressure delivered by a feedback-controlled mechanical probe. (i) Plots of the normalized peak amplitude of Ca^2+^ responses in individual TRPV1^+^ DRG neurons to increasing pressure. (j) Plots of the normalized peak amplitude of Ca^2+^ responses from TRPV1^+^ sensory afferents to the series of indentation stimuli. The peak amplitude was in a good linear correlation with the intensity level of indentation stimuli delivered (red dashed line, r^2^ = 0.98; n = 7 neurons). The pressure level that induced half maximum Ca^2+^ responses in TRPV1^+^ sensory afferents was close to 18 g/mm^2^. In comparison, the pressure level that induced half maximum Ca^2+^ responses in low-threshold mechanoreceptors was below 5 g/mm^2^. (k) Plots of the normalized decay time of Ca^2+^ responses (85%-15%) in TRPV1^+^ sensory afferents to the series of indentation stimuli. The decay time was in a good linear correlation with the intensity level of indentation stimuli delivered (r^2^ = 0.85; n=7 neurons).

We also analyzed whether the pinch-sensitive and –insensitive TRPV1^+^ thermoreceptors differ in morphology or response profiles. First, we found no difference in soma size between these two subgroups (Supplementary Fig. 4a; n = 18-40 neurons, p = 0.58). And although the average duration of the responses to 50°C was slightly longer in pinch-sensitive neurons (Supplementary Fig. 4c; n = 68-75 neuron, p = 0.016), the peak amplitude and decay were not significantly different between them (Supplementary Fig. 4b, d; n = 66-75 neurons, p = 0.08 for peak amplitude and p = 0.41 for decay).

### Individual TRPV1^+^ neuron encodes intensity in the noxious mechanical range

We next tested whether individual TRPV1^+^ neurons encode the intensity of mechanical indentation stimulus using a feedback-controlled mechanical probe. Stronger indentation stimuli induced larger responses in individual TRPV1^+^ neurons (Fig. 3h, i). The mean peak amplitude and decay time of Ca^2+^ responses were linearly correlated with the pressure intensity of the indentation stimuli (Fig. 3j, k). The pressure intensity that induced half maximum activity was 18 g/mm^2^ (Fig. 3j), which is in the range of pressure levels inducing nocifensive withdrawal responses using von Frey test in mice ^33^. Moreover, their lowest activation threshold was > 7 g/mm^2^. In contrast, low-threshold mechanosensory afferents, such as medium and large diameter TLR5^+^ neurons already displayed robust responses at 1.7 g/mm^2^, the lowest pressure intensity we tested (Fig. 3j). These results indicated that individual TRPV1^+^ neurons encode the intensity of noxious, but not innocuous mechanical stimuli.

The above results clearly indicated that TRPV1^+^ neurons should be involved not only in noxious thermoreception, but also in noxious mechanoreception, contrary to accepted thoughts ^13–15^. This finding also argues against a pure labeled line encoding scheme ^17–19,27^.

### Most MrgprD^+^ neurons are ploymodal nociceptors

Given our unexpected finding of polymodal TRPV1^+^ neurons, we decided to examine the sensory responses of MrgprD^+^ DRG neurons, which are thought to represent the counterparts of TRPV1^+^ afferents, i.e. to be the selective mechanoreceptive afferents. In virally transduced heterozygous MrgprD^Cre/+^ mice ^34^, > 85% of GCaMP^+^ neurons were MrgprD^+^ (Fig. 4a), indicating that most GCaMP labeled neurons expressed MrgprD. Moreover, most (88%) GCaMP^+^ neurons were IB4^+^, but only a small subset of GCaMP^+^ neurons were CGRP^+^ (15%), TRPV1^+^ (15%), or NF200^+^ (8%; Fig. 4b and Supplementary Fig. 5), demonstrating that the GCaMP^+^ neurons in MrgprD^Cre/+^ mice indeed represent non-peptidergic small-diameter DRG neurons ^35,36^.

**Figure 4.**
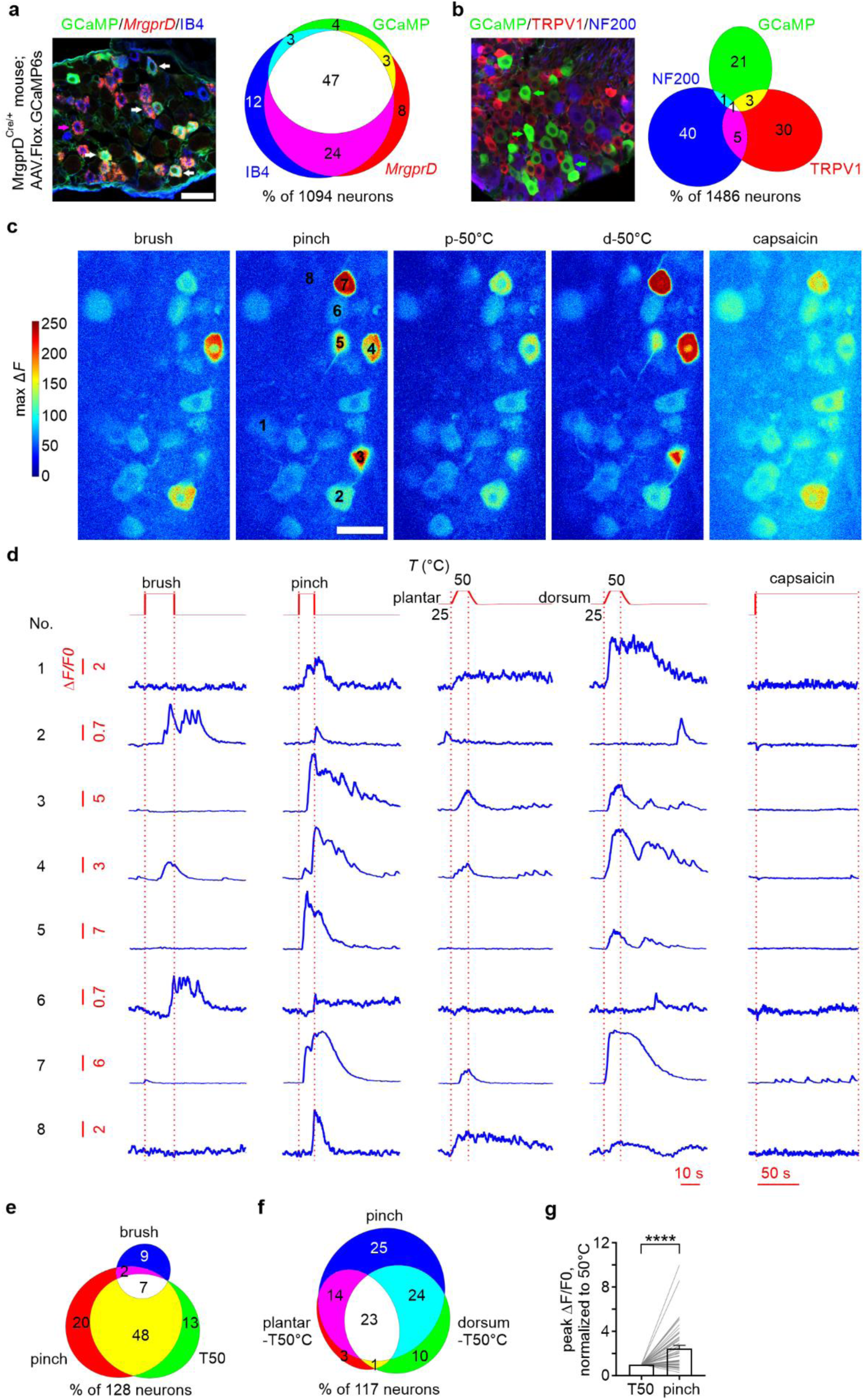
*In vivo* functional imaging of MrgprD^+^ sensory afferents. (a) Representative micrographs of DRG sections from MrgprD^Cre/+^ mice injected with AAV showing GCaMP6s expression (with anti-GFP antibody amplification) superimposed on labeling of *MrgprD* mRNA (RNAScope) and non-peptidergic neuron marker, IB4. The white arrows pointed to the neurons which were triple-positive. Euler diagrams showed that most GCaMP^+^ neurons expressed *MrgprD*. Scale bar = 50 µm. (b) Representative micrographs of DRG sections showing GCaMP6s expression (without antibody amplification) superimposed on labeling of TRPV1 and myelinated neuron marker, NF200. The green arrows pointed to the neurons which were only GCaMP6s positive. The cyan arrow pointed to the neurons which were GCaMP6s and NF200-double positive. Euler diagrams showed that most GCaMP^+^ neurons are unmyelinated and do not express TRPV1. (c) Color-coded heatmaps of a typical imaging field showing neuronal responses to mechanical and noxious heat stimuli in MrgprD^Cre/+^; AAV.Flex.GCaMP mouse. The noxious heat stimulation was applied to both plantar (p-T50°C) and dorsal side (d-T50°C) of hind paw in separate trials. Color scale indicates maximum △*F*. Scale bar = 50 µm. (d) Representative Ca^2+^ curves from eight neurons, highlighted in C, showing typical responses to mechanical and noxious heat stimuli. All eight neurons, except neuron 7, were insensitive to capsaicin stimulation. Note different Δ*F*/*F0* scales for the different neurons. The upper panels show the onset and offset of mechanical stimulation, the protocols for fast-ramp-and-hold thermal stimuli, as well as the bath application of capsaicin. Multiple repeated brushing or pinching stimulation was applied in each trial. (e) Euler diagram showing the overlap in neurons responsive to brushing, pinching, and noxious heat stimuli. Numbers are the percentage of the neurons responding to at least one of the three stimuli as indicated. (f) Euler diagram showing overlap in neurons responsive to noxious heat stimuli applied to the dorsal and plantar sides of the hind paw, as well as to pinching stimulation applied to both sides. Numbers indicated the percentage of the neurons responding to at least one of the three stimuli as indicated. (g) MrgprD^+^ neurons displayed significantly larger responses to pinching than noxious heat stimulation (n = 48 cells, ***p < 0.001 using Wilcoxon matched-pairs *t* test).

Consistent with being nociceptors, most MrgprD^+^ neurons responded to noxious pinching stimuli, applied to the hind paw (Fig. 4c-e). In contrast to TRPV1^+^ neurons, a small subset (18%) of MrgprD^+^ neurons also responded to gentle brushing of the skin (Fig. 4c-e). Surprisingly, however, the majority of MrgprD^+^ neurons also displayed robust responses to noxious heat (50°C) stimuli delivered to both sides of the hind paw (Fig. 4c-f). To confirm separation from TRPV1^+^ neurons, we applied capsaicin to the surgical opening at the end of the imaging session. More than 90% of GCaMP^+^ neurons were insensitive to capsaicin (Fig. 4c, d), consistent with the immunostaining results revealing that only a small subset (15%) of GCaMP^+^ neurons were TRPV1^+^ (Fig. 4b). Only capsaicin-insensitive neurons were included in the analysis.

Among all the capsaicin-insensitive MrgprD^+^ neurons that responded to at least one type of stimulus, 77% were pinch-sensitive, 18% were brush-sensitive, and 68% were 50°C-sensitive (Fig. 4e). Among pinch-sensitive neurons, 71% responded to 50°C stimulation (Fig. 4f). Slightly more pinch-sensitive neurons responded to the noxious heat stimulation applied to the dorsal side than the plantar side of the hind paw (Fig. 4f). On the other hand, more than 80% of noxious heat-sensitive neurons also responded to pinch stimulation (Fig. 4f), demonstrating that the majority of MrgprD^+^ neurons are polymodal nociceptors. Similar to TRPV1^+^ neurons, the responses of MrgprD^+^ neurons to pinch were also significantly larger than that to noxious heat (Fig. 4g).

### Thermosensitive MrgprD^+^ neurons are mostly selective for noxious heat

To further explore the thermosensitivity of MrgprD^+^ neurons, we applied the complete set of thermal stimulation to the plantar side of the hind paw (Fig. 5a, b). Among the MrgprD^+^ thermoreceptors, less than 20% responded to noxious cold (6°C) stimulation, while virtually no neurons responded to cold (15°C) and innocuous cool (20°C) stimulation (Fig. 5c). An increasing percentage of MrgprD^+^ thermoreceptors, responded to warm (30%), heat (50%), and noxious heat stimulation (90%). Virtually all MrgprD^+^ thermoreceptors were sensitive to 50°C stimulation (Fig. 5c).

**Figure 5.**
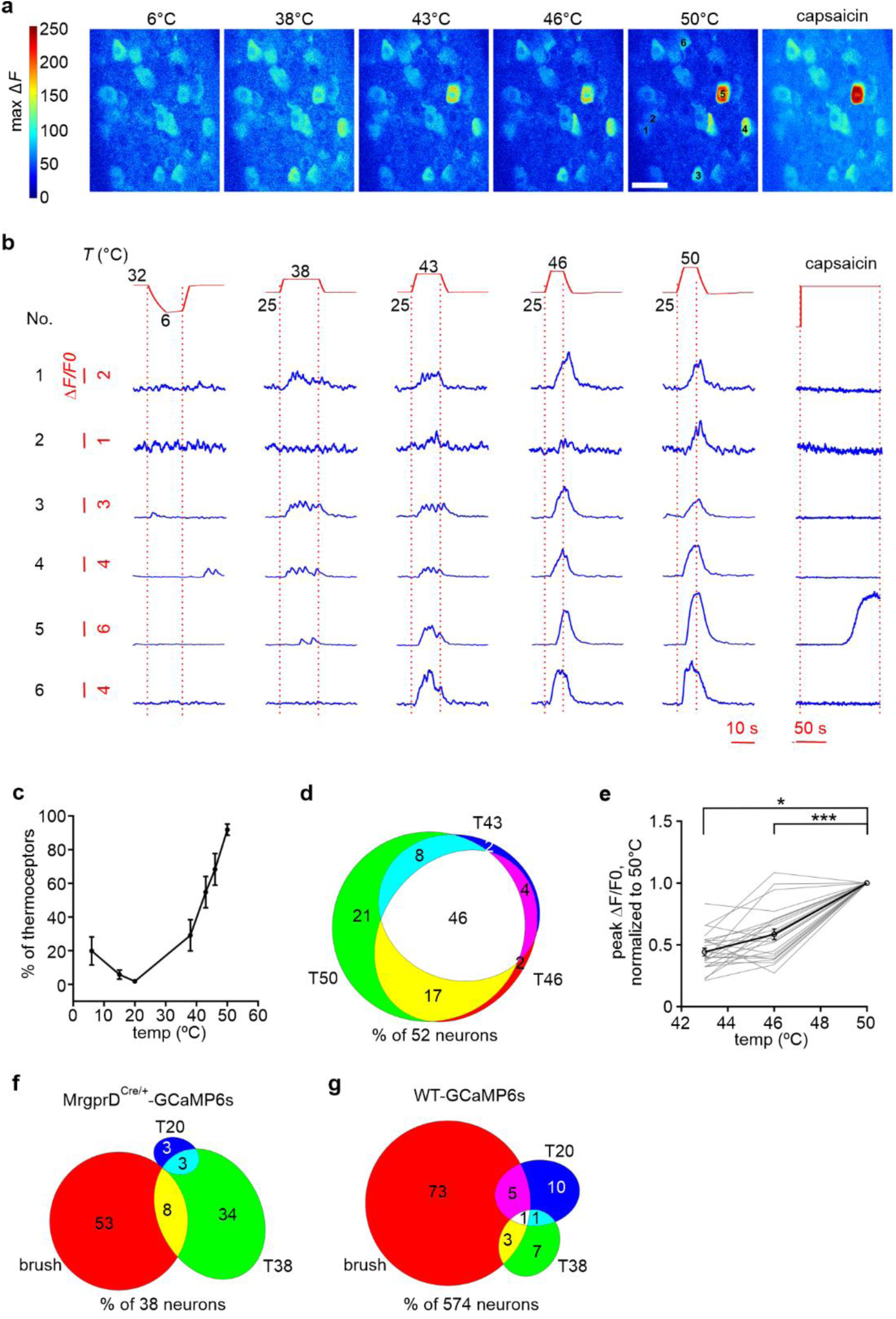
Most thermosensitive MrgprD ^+^ sensory afferents are heat nociceptors. (a) Color-coded heatmaps of a typical imaging field showing neuronal responses to innocuous and noxious thermal stimuli applied to the plantar side of the hind paw in MrgprD^Cre/+^; AAV.Flex.GCaMP mouse. Color scale indicates maximum △*F*. Scale bar = 50 µm. (b) Representative Ca^2+^ curves from six neurons, highlighted in A, showing typical responses to thermal and capsaicin stimulation. All six neurons, except neuron 5, were insensitive to capsaicin stimulation. Note different Δ*F*/*F0* scales for the different neurons. The upper panels show the protocols for fast-ramp-and-hold thermal stimuli applied to the plantar side of the hind paw, as well as the bath application of capsaicin. A baseline of 32°C and 25°C was chosen for cooling and heating stimuli, respectively. (c) Proportion of thermo-sensitive MrgprD ^+^ neurons (*i.e.* that responded to at least one type of thermal stimulus applied to the plantar side of the hind paw) in each imaging field that responded to different thermal stimuli (n = 13 fields from 13 mice). (d) Euler diagram showing the overlap in neurons responsive to the heat/noxious heat stimuli. Numbers indicated the percentage of the neurons responding to at least one of the three stimuli as indicated. (e) Plots of the normalized peak amplitude of “wide-range” heat-sensitive MrgprD^+^ neurons (gray curves = individual neuron response; black curve = mean response of the group) to increasing temperature. “Wide-range” heat neurons displayed significantly greater responses as temperature increased (n = 24 cells, *p < 0.05 and ***p < 0.001 using One-way Repeated Measures ANOVA test followed by Tukey’s multiple comparisons test). (f) Euler diagram showing the small overlap in MrgprD+ neurons responsive to brushing, innocuous warm, and innocuous cool stimuli. Numbers are the percentage of the neurons responding to at least one of the three stimuli as indicated. (g) Euler diagram showing the limited overlap in all types of DRG neurons responsive to brushing, innocuous warm, and innocuous cool stimuli. Numbers are the percentage of the neurons responding to at least one of the three stimuli as indicated. The data was recorded from wild-type mice.

Similar to TRPV1^+^ neurons, higher noxious temperature recruited more MrgprD^+^ neurons (Fig. 5d) and induced larger calcium responses in individual neurons (Fig. 5e), indicating that at both population and individual neuron levels, MrgprD^+^ neurons also encode the intensity of noxious heat.

### In the innocuous range, thermoreceptive and mechanoreceptive afferents are largely separated

Our functional imaging data revealed that about half of TRPV1^+^ and MrgprD^+^ afferents are polymodal, indicating that from the sensitivity perspective, thermo- and mechano-perception overlap extensively in the noxious range. Then we asked whether they also overlap in the innocuous range.

Although 20% of TRPV1^+^ afferents were innocuous warm-sensitive, none of them responded to innocuous cool or innocuous brushing stimuli (Fig. 2a, 3d). Small subsets (20 %) of MrgprD^+^ neurons responded to innocuous brushing or innocuous warm stimuli (Fig. 4e, 5c), but most (85%) of them responded to only one modality (Fig. 5f). Moreover, the data set we acquired from wild-type mice in a previous study ^25^, also showed that among all types of sensory neurons, the overlap between innocuous thermal-sensitive and innocuous mechanical-sensitive afferents was minimal (< 10 %; Fig. 5g). Thus, in sharp contrast to the noxious regime, where the level of cross-modality is >50%, primary afferents appear to encode innocuous thermal and mechanical inputs separately.

### TRPV1^+^ and MrgprD^+^ afferents are both necessary for thermo- and mechano-nociception

Given that a significant proportion of TRPV1^+^ afferents responded to both noxious thermal and mechanical input, we sought to test for the necessity of these afferents in thermo- and mechano-nociception. To selectively silence TRPV1^+^ neurons, we used low-concentration (0.05%) of capsaicin, which opens TRPV1 channel, to selectively permeabilize the cells to the membrane-impermeable quaternary derivative of lidocaine, QX-314, as previously described ^37,38^. Co-injection of QX-314 and capsaicin, but not QX-314 alone, or capsaicin alone, into the plantar area of hind paws, virtually completely blocked thermal nociception in both Hargreaves (Fig. 6a) and hot plate test (Fig. 6b). Consistent with our imaging data, the procedure significantly attenuated mechanical nociception, in both von Frey (Fig. 6c) and pin-prick (from 89% to 57% withdrawal responses; Fig. 6d) tests. Consistent with previous reports ^39^, our data indicate that TRPV1^+^ neurons are necessary for encoding not only thermal, but also mechanical nociception.

**Figure 6.**
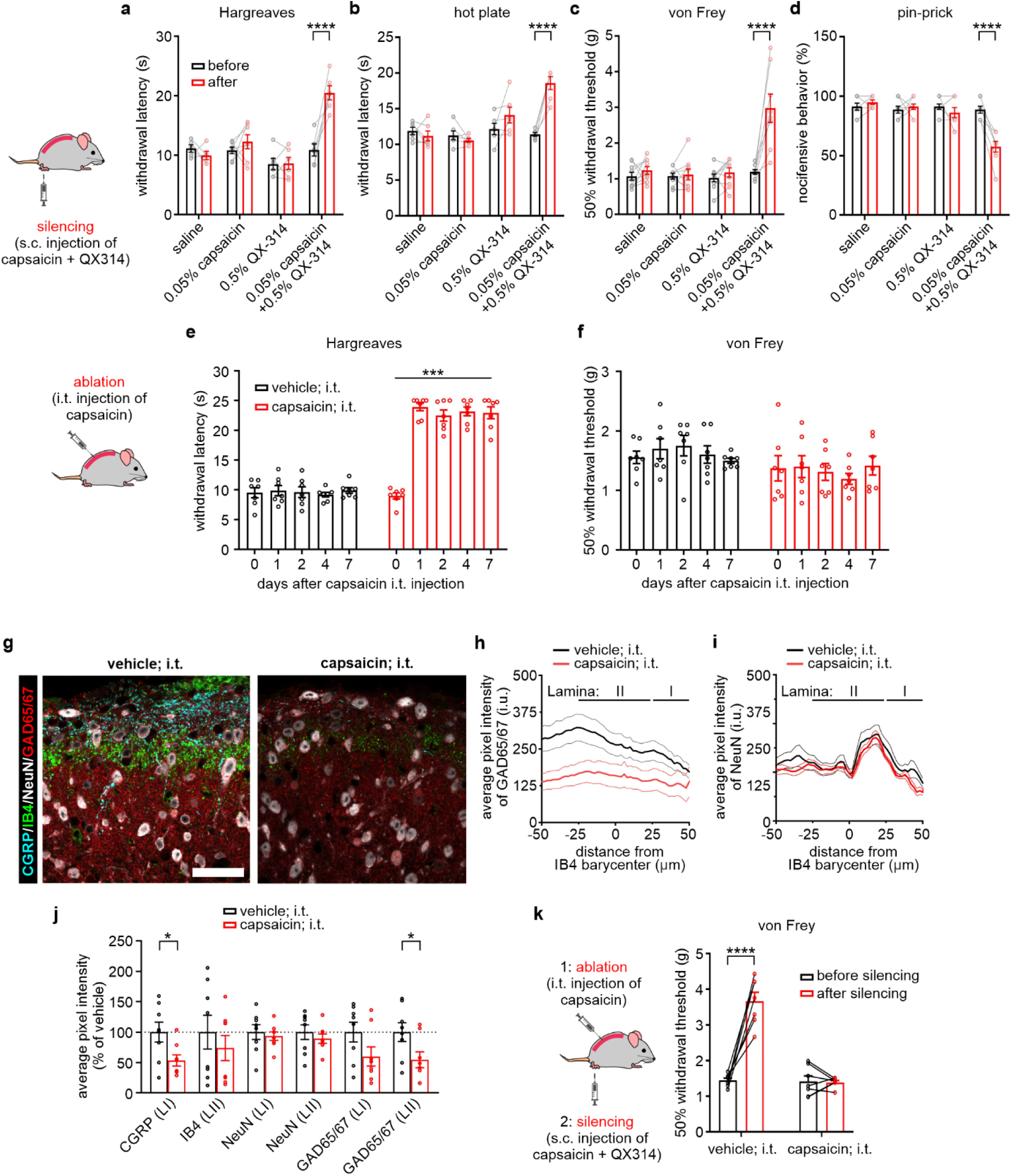
TRPV1^+^ sensory afferents are necessary for thermal and mechanical nociception. (a-b) Hargreaves (a) and hot plate (b) tests showed that silencing TRPV1^+^ neurons by combining capsaicin with QX-314 (membrane-impermeable sodium channel blocker), but not capsaicin alone, or QX-314 alone, profoundly blocked thermal nociception (n = 5-7 mice, ****p < 0.0001 using two-way Repeated Measures ANOVA test followed with Sidak’s multiple comparisons test). (c-d) Von Frey (c) and pin-prick (d) tests showed that silencing TRPV1^+^ neurons by combining capsaicin with QX-314, but not capsaicin alone, or QX-314 alone, also strongly inhibited mechanical nociception (n = 8-9 mice, ****p < 0.0001 using two-way Repeated Measures ANOVA test followed with Sidak’s multiple comparisons test). (e-f) Behavioral test of mice after intrathecal injection (i.t.) of vehicle or a high dose of capsaicin. The responses to noxious heat stimulation (Hargreaves test; e) of mice injected with capsaicin were completely abolished, while the responses to mechanical stimulation (von Frey test; f) were not affected (n = 7 mice for both vehicle and capsaicin groups, ***p < 0.001 using two-way Repeated Measures ANOVA test). (g) Representative confocal images of the spinal cord dorsal horn from mice injected intrathecally with either vehicle or capsaicin showing staining of CGRP, IB4, NeuN, and GAD65/67. Scale bar = 33.5 µm. (h) Average pixel intensities of GAD65/67 in the spinal cord dorsal horn as a function of the position towards the barycenter of IB4 labeling in vehicle i.t. injected (n = 8) *vs.* capsaicin i.t. injected mice (n = 8). (i) Average pixel intensities of NeuN in the spinal cord dorsal horn as a function of the position towards the barycenter of IB4 labeling in vehicle i.t. injected (n = 8) *vs.* capsaicin i.t. injected mice (n = 8). (j) Average pixel intensity in vehicle and capsaicin injected i.t. animals, of CGRP in lamina I, of IB4 in lamina II, of NeuN, and of GAD65/67 in lamina I and II of the spinal cord dorsal horn (vehicle n = 8 mice; capsaicin n = 8 mice; *p < 0.05). (k) Von Frey tests showed that combining capsaicin with QX-314 did not have an analgesic effect on the mice in which TRPV1^+^ afferents were ablated (n = 7 mice, ****p < 0.0001 using two-way Repeated Measures ANOVA test followed with Sidak’s multiple comparisons test).

These findings, however, seem to be in contradiction with previous reports where ablation of TRPV1^+^ afferents appeared to selectively abolish thermal nociception, with no effect on mechanical nociception ^13–15^. To test whether this discrepancy may be due to the difference in the procedures for silencing, we repeated their experiments using intrathecal (i.t.) injection of high concentration (0.2%) of capsaicin into adult wild-type mice to ablate TRPV1^+^ afferents as reported^13^. One to seven days after injection, behavioral tests showed that ablating TRPV1^+^ afferents abolished thermal nociception (Fig. 6e), but left mechanical nociception intact (Fig. 6f), in concordance with previous findings ^13–15^, but contradicting our findings with QX314-mediated silencing and functional imaging data. We reasoned that the strong neuronal activity and neuropeptide release by i.t. capsaicin administration at the spinal level might have additional effects other than ablation of TRPV1^+^ afferents ^40,41^. To test this hypothesis, we assessed the state of spinal inhibitory circuits. First, we examined the expression of glutamic acid decarboxylase 65 (GAD65) and GAD67 isoforms, which synthesize GABA, the main inhibitory neurotransmitter ^42–44^. Quantitative immunocytochemical analysis using an anti-GAD65/GAD67 antibody ^45^ revealed a significant decrease in expression in the superficial dorsal horn seven days after i.t. capsaicin-induced ablation (Fig. 6g, h, j, and Supplementary Fig. 6a). While the marginal decrease in inhibitory terminals seen in spinal lamina I may reflect the loss of presynaptic boutons due to the ablation of TRPV1^+^ and CGRP^+^ afferents terminals (Fig. 6g, j), we also found a significant decrease in the numbers of inhibitory terminals in lamina II (Fig. 6g, h, j, and Supplementary Fig. 6a). This indicated that the effect of i.t. high-dose capsaicin was not spatially limited to lamina I where the TRPV1^+^ afferents mainly innervate.

As a second proxy of inhibitory capacity, we analyzed the expression of the neuron-specific K^+^-Cl^−^-cotransporter, KCC2 ^45–47^. Its activity maintains the low concentration of intracellular Cl^−^, which is critical to maintain the efficacy of glycine and GABAA-mediated inhibition in mature central nervous system neurons ^46,48,49^. In contrast to GAD65/67, the expression of KCC2 was stable after i.t. capsaicin-mediated ablation (Supplementary Fig. 6b, c), indicating that capsaicin does not alter the capability of superficial spinal cord neurons to be inhibited by GABA and glycine ^49–52^. As a negative control for both GAD isoforms and KCC2 immunostaining, the staining intensity of an anti-NeuN antibody was stable, suggesting the number of spinal cord neurons was unchanged (Fig. 6i, j). Collectively, our histological data revealed that there was a loss of inhibition in the spinal cord in addition to ablation of TRPV1^+^ afferents upon the i.t. administration of the high dose of capsaicin.

Importantly, co-injection of QX-314 and capsaicin did not affect noxious mechanosensitivity in mice whose TRPV1^+^ afferents were already ablated (Fig. 6k), further confirming that QX-314 and capsaicin selectively affect TRPV1^+^ neurons.

Our imaging results showed that a majority of MrgprD^+^ neurons are polymodal nociceptors. To test their role in nociception, we used a chemogenetics approach ^53^ to acutely inhibit MrgprD^+^ neurons in behavioral experiments. First, we used viral injection to express inhibitory DREADDs (hM4d) in MrgprD^+^ neurons. Then, intraperitoneal injection of CNO, but not saline, significantly attenuated mechanical nociception, in both von Frey (Fig. 7a) and pin-prick (Fig. 7b) tests, and also reduced thermal nociception in both Hargreaves (Fig. 7c) and hot plate (Fig. 7d) tests. No effect of CNO was observed on the MrgprD-Cre mice transduced with a control virus (AAV2/9-CAG-DIO-mCherry). Collectively, our results revealed that both MrgprD^+^ and TRPV1^+^ primary afferents are each involved in both mechanical and thermal nociception. In contrast, we found that <10% overlap in thermal and mechanical modalities in the innocuous range among all types of afferents (Fig. 5g). This indicates that thermal and mechanical sensitivity converge only in the nociceptive range.

**Figure 7.**
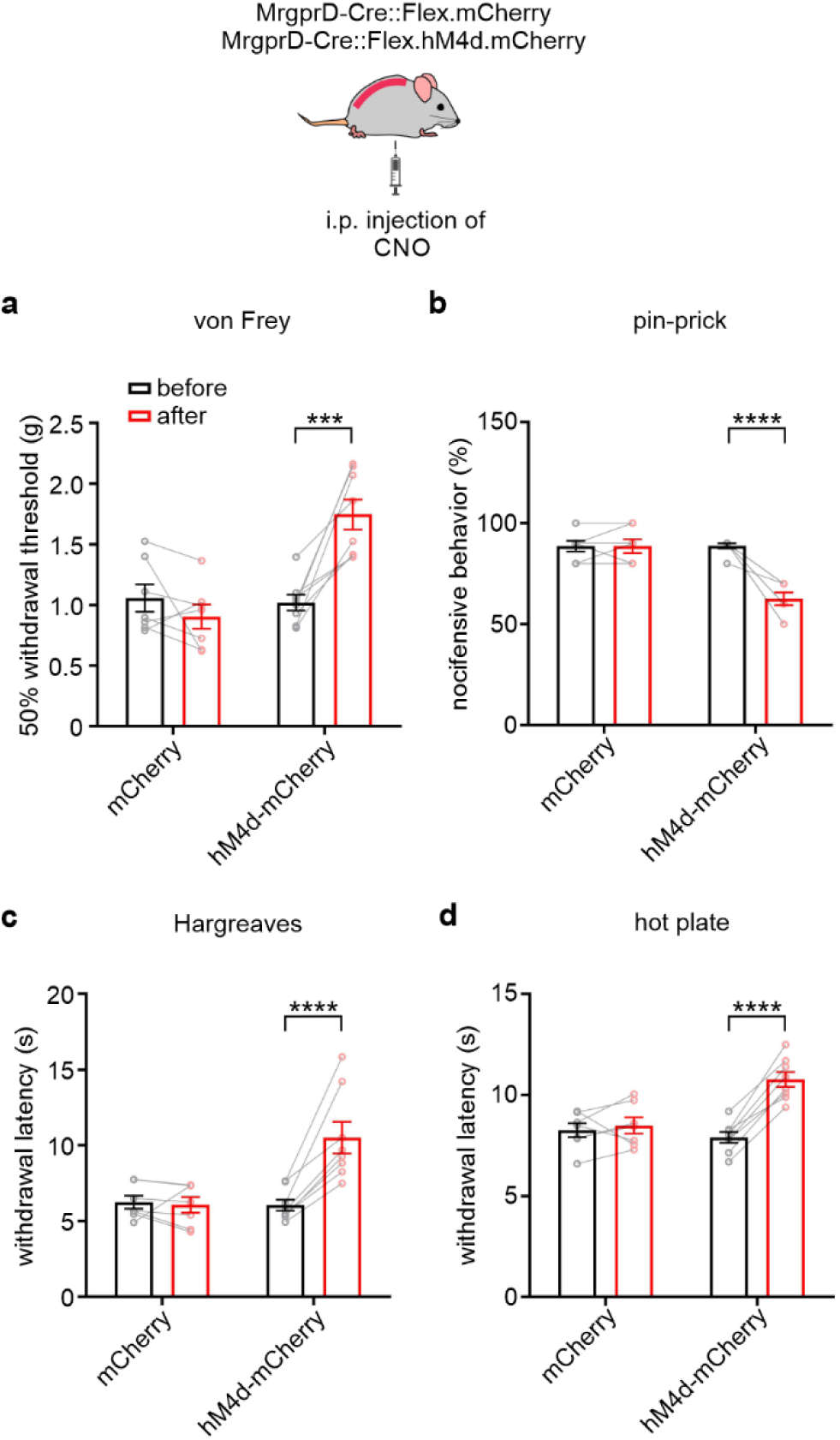
MrgprD^+^ afferents are essential for thermal and mechanical nociception. (a-b) Inhibiting MrgprD^+^ neurons with inhibitory DREADDs attenuated mechanical nociception in both von Frey (a) and pin-prick (b) tests (n = 7-8 mice, both ***p and ****p < 0.0001 using two-way Repeated Measures ANOVA test followed with Sidak’s multiple comparisons test). (c-d) Inhibiting MrgprD^+^ neurons with inhibitory DREADDs also reduced thermal nociception in both Hargreaves (c) and hot plate (d) tests (n = 7-8 mice, both ***p and ****p < 0.0001 using two-way Repeated Measures ANOVA test followed with Sidak’s multiple comparisons test).

## Discussion

By recording from large populations of genetically defined primary sensory neurons in live animals, we comprehensively assessed the representation and the strategies of nociception coding at the peripheral. Unexpectedly, our results revealed that the majority of TRPV1^+^ and MrgprD^+^ neurons are polymodal nociceptors. Consistently, behavioral experiments using acute neuronal blockade demonstrated that the two subpopulations of nociceptors are necessary for both mechano- and thermo-nociception. In contrast, in the innocuous regime, mechano- and thermoreception appear largely separate among afferents, suggesting that different modalities only merge in the noxious range.

### Thermal and mechanical sensitivity of TRPV1^+^ and MrgprD^+^ neurons

It has been long suggested that TRPV1^+^ and MrgprD^+^ primary sensory neurons are well-separated labeled lines for heat- and mechano-nociception, respectively, based on their distinct innervation patterns in the skin and spinal cord ^36,54^, as well as the results from behavioral studies, but mainly using ablation approaches ^13,14^. However, the physiological sensitivity of these two subpopulations of sensory neurons remains largely unclear, especially under *in vivo* conditions. Our Ca^2+^ imaging approach revealed that *in vivo* about half of TRPV1^+^ and MrgprD^+^ afferents are polymodal nociceptors (Fig. 3d, 3e, 4e, 4f), consistent with previous studies in which non-peptidergic IB4^+^ and MrgprD^+^ neurons were found to be polymodal in an *ex vivo* (skin/nerve/DRG/spinal cord) preparation ^21,34,55^. Moreover, a very recent study using *in vivo* imaging approach also revealed that a subpopulation of MrgprD^+^ neurons are high-threshold polymodal nociceptors ^56^. In contrast, patch recording from DRG neurons in the *ex vivo* preparation, combined with post-hoc labeling, showed that TRPV1^+^ neurons were noxious heat-specific and mechano-insensitive ^21^. However, it is worth noting that, our *in vivo* imaging approach better preserves the biological integrity of the periphery system and remains closer to the physiological conditions. It will be intriguing to study how the TRPV1^+^ neurons lose their mechano-sensitivity in the *ex vivo* preparation.

### The central circuits of TRPV1^+^ and MrgprD^+^ afferents

In the mice spinal cord, TRPV1^+^ afferents innervate lamina I and outer lamina II, while MrgprD^+^ afferents innervate inner lamina II, which appears to be segregated pathways ^54^. Furthermore, some spinal cord interneurons only receive exclusive input from one of these two subpopulations. For example, it has been reported recently that heat-selective ErbB4^+^ spinal interneurons are only innervated by TRPV1^+^ afferents, but not MrgprD^+^ afferents ^57^. However, similar to nociceptors, the majority of dorsal horn interneurons and a substantial subpopulation of spinal projection neurons are also polymodal ^58–61^, indicating that the information relayed by these two subpopulations of primary afferents can eventually converge to spinal projection neurons. Consistently, at the anatomical level, polymodal nociceptive projection neurons in lamina I receive dense direct innervation from peptidergic DRG neurons ^62^, which largely overlap with TRPV1^+^ neurons, and also indirect input from MrgprD^+^ afferents, probably via excitatory vertical interneurons ^63,64^, which are located in outer lamina II. However, the indirect input from MrgprD^+^ afferents to projection neurons appears to be under strong inhibitory control as stimulating only TRPV1^+^ afferents, but not MrgprD^+^ afferents, activated superficial projection neurons ^65^. Overall, the interaction and integration of the information carried by TRPV1^+^ and MrgprD^+^ afferents in the spinal cord remains poorly understood.

### TRPV1^+^ and MrgprD^+^ afferents are essential for both mechanical and thermal nociception

In our study, acute silencing of TRPV1^+^ sensory neurons almost completely blocked both thermal and mechanical nociception and pharmacogenetic inhibition of MrgprD^+^ neurons also resulted in deficits of both, in contrast to the results of studies using ablation approaches ^13–15^. One possible explanation is that ablation strategies have the caveat of potentially triggering compensatory mechanisms in somatosensory circuits ^16^. Our finding that GAD65/67 labeling in lamina II was significantly reduced, indicating impairment of inhibition, confirmed this assertion. Similar discrepancies between the results of acute inhibition and ablation experiments have been reported in the spinal cord neurons as well. For example, genetic ablation of somatostatin^+^ lineage spinal cord neurons caused a profound and selective loss of mechanosensitivity ^66^, while pharmacogenetic inhibition of the same population of neurons resulted in deficits of both mechano- and thermo-nociception ^67^, also indicating possible long-term compensatory mechanisms following genetic ablation. On the other hand, our approach of acute silencing has better specificity with less potential for compensation. In agreement with our results, acute inhibition of TRPV1^+^ DRG neurons with an optogenetics approach significantly attenuated both thermal and mechanical nociception ^68^, demonstrating that TRPV1^+^ sensory neurons are indeed essential for both thermal and mechanical nociception.

With respect to the observation that pathological pain is categorised in terms of perceived location or etiology, not modality, a key question is whether activation of TRPV1^+^ and MrgprD^+^ DRG neurons induces the same sensation or not. It has been reported that activating either TRPV1^+^ or MrgprD^+^ DRG neurons with optogenetics approaches could induce nocifensive behavior in mice, but with subtle differences ^16^. Transdermal light stimulation of TRPV1^+^ neurons generated paw withdrawal and licking, while stimulation of MrgprD^+^ afferents generated mainly withdrawal and lifting ^16^. However, these two phenotypes can result from distinctions in intensity rather than modalities. Indeed, although activating both populations causes nociception, activation of TRPV1^+^ neurons likely induces stronger pain than activation of MrgprD^+^ neurons because licking behavior is often associated with stronger pain than lifting behavior ^69,70^. Our behavioral results also largely agree with this interpretation, as silencing TRPV1^+^ afferents induced stronger analgesia than MrgprD^+^ neurons (Fig. 6a-d and 7). Moreover, only activation of TRPV1^+^ neurons, not MrgprD^+^ neurons, induces conditioned place aversion ^16,55^, which may reflect that only TRPV1^+^ neuron activity is sufficient to yield significant aversion. Thus, current behavioral phenotypes in mice are insufficiently informative to assert discrimination on the basis of modality *vs.* intensity. In humans, capsaicin evokes a fast stinging sensation followed by burning pain ^71^, while β-alanine (ligand of MRGPRD receptor) evokes itching and stinging sensation as well as burning ^72^, indicating that activation of TRPV1^+^ and MrgprD^+^ afferents induces similar sensations.

### Thermoreceptive and mechanoreceptive afferents merge in the noxious range, while they are largely separated in the innocuous range

In mice DRGs, TRPV1^+^ and MrgprD^+^ are virtually separated subpopulations of sensory afferents^29^. In the innocuous range, our imaging data revealed that TRPV1^+^ afferents only responded to innocuous warm stimulation, but not to innocuous cool or gentle brushing stimulation (Fig. 2a, 3d). Although some MrgprD^+^ afferents are sensitive to brushing stimulation or warm stimulation, they are small subpopulations with limited overlap (Fig. 5f). In addition, data from our previous study using wild-type mice showed that the overlap among innocuous thermal and mechanosensitive primary afferents is also very limited (Fig. 5g). Consistently, although some low-threshold mechanoreceptors are sensitive to temperature decrease ^73^, the majority of them are different from nociceptive and thermosensitive afferents ^74^. Collectively, these findings suggested that thermoreceptive and mechanoreceptive afferents in mice are largely separated in the innocuous range, while they merge in the noxious range, as revealed by our imaging and behavioral results. The observation that when TRPV1^+^ afferents have some sensitivity in the innocuous range, it is only to warmth adds to the argument above that noxious heat sensation is difficult to dissociate from contextual innocuous warm sensation, which may underlie the association of nociception to the *thermal* modality when a thermal stimulus is triggering it.

The degree of convergency in the noxious range might be even higher in other species. Anatomically, the mouse species presents an extreme case of separation of peptidergic vs non-peptidergic, or restriction of TRPV1 to a subgroup of nociceptors ^29^. The overlap of peptidergic and non-peptidergic neurons is substantial in rats ^75–77^ and there is almost no such separation in humans as the majority of human DRG neurons were found to be cRet and TRPV1-positive ^78,79^. Single units recordings in monkeys ^80^ and microneurography studies in humans ^81,82^ also reported high levels of multimodality from C-nociceptors, responding to both thermal and mechanical noxious stimulations.

Our finding that thermal and mechanical modalities merge in the noxious range is consistent with the concept that, for survival, nociceptors may convey a threat signal that does not need to be associated with the originating modality.

## Methods

### Animals

All experiments were performed in accordance with regulations of the Canadian Council and Animal Care. In this study, Trpv1^Cre^ knock-in mice were purchased from The Jackson Laboratory (B6.129-*Trpv1^tm1(cre)Bbm^*/J; 017769). Gene-deleting *Mrgprd-Cre*^−/-^ mice were purchased from MMRRC (Mrgprd^tm1.1(cre)And^; 36118). Heterozygous mice produced by crossing homozygous TRPV1^Cre^ or *Mrgprd-Cre*^−/-^ mice with wild-type C57BL6/J mice (The Jackson Laboratory; 000664) were used in most experiments, while homozygous TRPV1^Cre^ mice were also used in the immunostaining experiment as specified. Male adult wild-type C57BL6/N mice were purchased from Charles River (strain 027) for behavioral tests. Animals were maintained on a 12:12 hour light:dark cycle with food and water provided ad libitum.

### Virus injection

Newborn mice pups at P3 to P7 were used for virus injection. Before the injection, the mother was anesthetized with low concentration of isoflurane (1%) and pups were anesthetized by hypothermia. Virus AAV9.CAG.Flex.GCaMP6s.WPRE.SV40 (UPenn viral vector, AV-9-PV2833 or Addgene, 100842-AAV9), AAV9.CAG.DIO.hM4D(Gi).mCherry (Canadian Neurophotonics Platform Viral Vector Core Facility; construct-639), and AAV9.CAG.DIO.mCherry (Canadian Neurophotonics Platform Viral Vector Core Facility; construct-1448), was injected into the plantar area of the hind paw using a 50 µl Hamilton syringe equipped with a 30G x 1/2 needle. 5 µl of virus with titer above 1×10E12 GC/ml was injected into one hind paw, and both hind paws were injected. After injection, the pups were kept on a heating pad until they woke up and the body temperature returned to normal. Then they were put back into the cage before the mother was put back and rubbed against the pups. The mice injected with virus were separated from the mother after weaning and used for experiment 6 to 8 weeks after virus injection.

### Immunostaining

For immunohistochemistry, mice were transcardially perfused with saline and followed by 4% w/v paraformaldehyde and 0.1% v/v picric acid in phosphate buffer (in mM: 100 KH2PO4 and 80 NaOH; pH = 7.4). The DRGs and spinal cord were dissected and post-fixed for 2 h in the same fixative solution. The tissue was cryoprotected in 30% sucrose solution overnight.

DRGs were sectioned using a cryostat at 10 µm thickness. The sections were incubated with the primary antibodies (refer to Table 1 for the list of the primary antibodies used) at 4°C overnight. Then they were washed three times with PBS (in mM: 155.17 NaCl, 1.06 KH2PO4, and 3.00 Na2HPO4; pH = 7.4) and incubated with the secondary antibodies and/or IB4 dyes (refer to Table 2 for the list of the secondary antibodies and IB4 dyes used) at room temperature for three hours. DRG sections were examined under a confocal laser scanning microscope (Zeiss LSM700) using a 20x objective.

The fixed spinal cord tissue was sectioned transversally using a vibratome in 25 µm-thick slices. Sections were washed twice in PBS and incubated overnight at 4°C on a shaker with three types of primary antibody mixtures (refer to Table 1 for the list of the primary antibodies used) containing 10% normal goat serum in PBST (PBS containing 0.3% Triton X-100). On the second day, sections were washed three times in PBST, then sections were incubated at room temperature for 2 hours in a mixture of the secondary antibodies and IB4 dyes (refer to Table 2 for the list of the secondary antibodies and IB4 dyes used). Sections were washed three times in PBS before being mounted on Superfrost plus microscope slides (Fisherbrand catalog No. 12-550-15) and coverslipped using fluorescence mounting medium containing DAPI dye (catalog #S3023, Dako).

Spinal cord images were captured using a confocal laser scanning microscope (Zeiss, 880). Appropriate fluorescent probes excitation parameters were selected for the separate detection of DAPI, Alexa Fluor 488, Cy3, and Alexa Fluor 647, using a multitrack scanning method. Twelve-bit images were taken with a 63x/1.4 oil-immersion objective lens at a resolution of 2048 x 2048 pixels with a pixel dwell time of 1.02 µs and a pixel size of 0.066 µm. To ensure consistency among samples, all parameters of laser power, pinhole size, and image detection were kept unchanged between the image acquisitions of different samples. The chosen parameters were set so that the detection of the staining was maximal while avoiding pixel saturation.

### Inter-laminar profile expression of GAD65/67, neuronal membrane KCC2 and NeuN, in the superficial dorsal horn (SDH)

To determine inter-laminar variations of the inhibitory terminals and membrane KCC2 in the SDH after intrathecal injection of capsaicin (see below), we measured their distribution by immunohistochemistry. As described in the adult C57BL/6 mouse, we used CGRP, IB4, and NeuN to delineate SDH laminae ^52,83^. To avoid biased quantification due to differences in the laminar size between sections, GAD65/GAD67^+^ terminals, KCC2 at the neuronal membrane, and NeuN profile intensities were expressed as a function of the distance from the IB4 barycentre ^52^. Intensity profiles of NeuN were used as a control for the other markers.

### In situ hybridization

*In situ* hybridization was carried out using RNAScope kit from Advanced Cell Diagnostics (ACD). Briefly, DRG sections were treated with Proteinase Plus (322330, ACD) for 10 minutes at 40°C. Then, the sections were incubated with *MrgprD* (417921, ACD) probes for two hours at 40°C. Afterwards, the signal was revealed using RNAscope 2.5 HD Detection Reagents-Red (322360, ACD).

The GCaMP6s signal was not detectable after *in situ* hybridization, so immunostaining using anti-GFP antibodies was applied to reveal GCaMP6s expression. Immediately after *in situ* hybridization, DRG sections were washed three times with PBST (in mM: 81 Na2HPO4, 21.9 NaH2PO4, 154 NaCl; 0.15% Triton X-100, pH = 7.2-7.4) and then incubated with a mouse anti-GFP (1:500; A-11120, Invitrogen) diluted in PBST containing 4% normal goat serum at 4°C for 48 hours, followed by incubation with an Alexa Fluor 488-conjugated goat anti-mouse secondary antibody (1:100; Molecular Probes) at 37°C for two hours.

### Single-cell PCR

Dissociated DRG neurons were prepared according to the previous report ^84^. Single DRG neurons were randomly sucked into a glass pipette, and gently transferred into lysis buffer. Reverse transcription was performed using a SMARTer Ultra Low RNA Kit (Clontech) in the cell lysate while single-cell cDNA was amplified using an Advantage 2 PCR Kit (Clontech) according to the manufacturer’s protocol. The amplified cDNA was kept for single-cell PCR experiments. PCR was performed with Ex Taq (Takara). The expression was detected using following primers: 5′ GCCTCTGTCGCCTCCAAA 3′ and 5′ CAGGTGCCCAGCAGTGAA 3′ (Piezo2); 5′ GCGAGTTCAAAGACCCAGAG 3′ and 5′ACATCTGCTCCATTCTCCAC3′(Trpv1).

### Laminectomy

Six weeks after virus injection, the mice were deeply anesthetized intraperitoneally with 100 mg ketamine, 15 mg xylazine, and 2.5 mg acepromazine (A7111, Sigma-Aldrich) per kg in saline ^85,86^. Animals were kept anesthetized for the duration of the *in vivo* imaging experiments with repeated hourly injections of one-fourth of the original dose. The back of the animals was shaved and a midline incision of the skin exposed the back musculature over the lumbar vertebrae. The paravertebral muscles at the desired level (L6-S1) were carefully separated from the vertebral column (Harrison *et al.*, 2013). Laminectomy of a single vertebra was performed to expose L4 DRG. Then the spinal columns flanking the laminectomy exposure were clamped with two clamps of a home-made spinal stabilization device to fix the animals. 3% agarose solution was used to make a pool for holding Ringer solution (in mM: 126 NaCl, 2.5 KCl, 2 CaCl2, 2 MgCl2, 10 D-Glucose, and 10 HEPES; pH = 7.0). To label the blood vessels, mice were injected intravenously with 1% dextran Texas Red (70 kDa, Neutral, D1830, Invitrogen) in saline. The animal was heated with a heating pad during the surgery and imaging to keep the body temperature at 37℃. Warmed Ringer solution was dropped on the exposed spinal cord and DRG to keep the moisture, and Ringer was changed from now and then during the imaging experiment.

### In vivo Ca^2+^ imaging

Animals with the whole spinal stabilization device were fixed on a home-made video-rate two-photon microscope ^87^. A tunable femtosecond laser (Spectra-Physics, InSight X3) was set to 940 nm for imaging. Filters for wavelength selection between 500-550 nm and above 605 nm light were used for GCaMP6s and Texas Red, respectively. A water-immersion 40x objective (Olympus) was used and the resolution of the raw images was 0.375 µm/pixel. Images were acquired at 32 frames/second. For thermal stimulation, a feedback-controlled Peltier device and a 1 cm by 1 cm thermal probe (TSA-Ⅱ -NeuroSensory Analyzer, Medoc) were used to deliver different temperatures, ranging from 6°C to 50°C, in a fast ramp and hold mode to the plantar side of hind paws. The thermal stimuli were triggered by the acquisition software using the data acquisition card (LabJack, U3-HV). The speed of temperature increase was set at 8°C/s, and temperature decrease at 4°C/s. The steady-state duration of the 38°C, 43°C, 46°C, and 50°C stimuli was 15 s, 10 s, 5 s, and 5 s, respectively. The baseline temperature for heating stimuli was set to 25°C. For cold stimulation, the baseline temperature was set at 32°C, and the duration of the steady-state phase of the 20°C, 15°C, and 6°C stimuli was 15 s, 10 s, and 8 s, respectively. The relatively shorter duration of noxious heat and cold stimuli was chosen to avoid sensitization and injury. Noxious heat (50°C) was also tested on the dorsal side of the hind paws.

An innocuous brushing on the hind paw was performed using a small painting brush. Noxious mild pinching at the hind paws was applied using serrated forceps. Mechanical stimuli (brushing and pinching) were applied multiple times during one stimulation epoch, which was 15 s (brushing) or 8 s (pinching) long. There were 3 to 5 minutes intervals between any two stimulations to avoid neuronal sensitization.

### Mechanical probe

For mechanical stimulation, a dual-mode muscle lever (300C; Aurora Scientific) was used to deliver a series of indentation stimuli to the plantar area of the hind paw. The force delivered was calibrated with a force gauge. A round tip with a diameter of 1 mm was equipped so the pressures of the indentation stimuli reached the noxious range. The mechanical probe was also triggered by the acquisition software using the same data acquisition card (LabJack, U3-HV).

### Ca^2+^ imaging data analysis

Acquired raw image sequences were converted to TIFF format in ImageJ. Then the TIFF movies were registered with a custom-built MATLAB (MathWorks) function through rigid body translation alignment based on the 2D cross-correlation to correct for movement. A rectangular region of interest (ROI) in a region without any visible neuron was drawn as background ROI. The average pixel value inside the background ROI for each frame was subtracted from every pixel in the corresponding frame to remove excess noise. Neuronal ROIs were drawn manually in the cytoplasm of visible neurons. The average fluorescence intensity of a given ROI, Ft, was measured by averaging pixel values inside the ROI. For Ca^2+^ traces, ΔF/F0 = (Ft-F0)/F0, where F0 is the fluorescence value at baseline, measured as the average of the first two seconds of Ft. To avoid aberrant amplification due to small F0 values in some neurons (*e.g.,* low basal fluorescence), when it was <1, F0 in the denominator, but not in the numerator, was set to 1. The resulting Ca^2+^ time series extracted from the image sequences were synchronized with thermal or mechanical stimulus data series. This part of the processing was performed using custom functions written in MATLAB.

Positive responses were detected and measured automatically using another custom tool written in Spike2 (CED), which provides an integrated interface to visualize and analyze traces. Raw Ca^2+^ traces were first smoothed with a 1 s temporal window. The baseline was selected from a period starting 1 s after the beginning of the recording and ending 1 s before the stimulus onset (usually 3 to 5 s in duration). Then, Fb and Fb-max were calculated as average and maximum ΔF/F0 values during baseline, respectively. A response was considered positive when the peak of the Ca^2+^ trace during stimulation was above Fb+(Fb-max-Fb)*x, where x was a value between 2 and 3, depending on baseline stability, to provide the most reliable detection. Given the relatively slow decay of GCaMP6s, responses with very brief duration (< 0.5 s) were excluded. While the detection algorithm was found to be highly reliable, all traces were also visually inspected to ensure no false positives were included and no false negatives were missed. Peak amplitude and decay time (from 85% to 15% of peak amplitude) were measured for all responses using automated algorithms. For each neuron, each parameter was measured for each of the positive responses obtained over multiple trials for each stimulus and then averaged. Finally, all the data was merged into a single database and Microsoft Excel Pivot Table was used to select the proper subsets for subsequent statistical and cluster analyses. Part of the imaging data from TRPV1^+^ neurons was published before ^25^.

### Behavior test

All animals were acclimatized to the behavioral testing apparatus in three to five ‘habituation’ sessions. Thermal sensitivity was assessed using Hargreaves and hot plate tests. Thermal sensitivity apparatus (IITC Life Sciences) was used for the Hargreaves test. Mice were placed within small (12 cm by 8 cm) plexiglass cubicles on a 3/16th-inch thick glass floor warmed to 30°C, and a focused high-intensity projector lamp beam was shone from below onto the mid-plantar surface of the hind paw. A 25-second cutoff was set, and the latency to withdraw from the stimulus was measured to the nearest 0.1 s. Both hind paws were tested three times under each condition and all results were averaged for each experimental condition.

For the hot plate test, mice were placed, one by one, on the hot plate (BIOCHP; Bioseb) maintained at 55°C, with their locomotion restricted by a plexiglass cylinder. The latency to hind paw licking and/or shaking, or jumping, was recorded, and then the animal was immediately removed from the hot plate. A cutoff of 20-seconds was set to avoid tissue injury. At each condition, all animals were tested sequentially twice with a minimum of 10 minute intervals, and the values were averaged.

Mechanosensitivity was measured using von Frey and pin-prick test. The classical up-down method with von Frey filaments was used to estimate the 50% withdrawal threshold in gram units. Both hind paws were tested under each condition and results were averaged for each experimental condition.

For the pin-prick test, we gently touched the plantar surface of the hind paw with a pin without skin penetration. Each hind paw was measured 5 times with 1-minute intervals and both hind paws were measured. The number of withdrawal responses was recorded.

### Intrathecal injection of capsaicin

To kill TRPV1^+^ afferents, 5 µl of high-concentration capsaicin (2 µg/µl) was injected intrathecally into wild-type C57BL6/N mice. The capsaicin was diluted in the vehicle, which was saline containing 10% ethanol (v/v) and 10% Tween 80 (v/v). The thermal and mechanical sensitivity of the injected mice were measured before, 1 day, 2 days, 4 days, and 7 days after the injection. Then the mice were perfused and the spinal cords were processed for histological studies.

### Acute blockage of TRPV1^+^ sensory afferents

The thermal and mechanical sensitivity of wild-type C57BL6/N mice were tested before injection as a control. Then mice were anesthetized with 1% isoflurane and 20 µl of saline, or 0.5% QX-314 (in saline), or 0.05% capsaicin (in saline), or 0.5% QX-314 plus 0.05% capsaicin was injected into the plantar area of both hind paws. Half an hour after injection, the same behavioral test was repeated.

### Chemogenetic inhibition of MrgprD^+^ sensory afferents

The thermal and mechanical sensitivity of MrgprD-hM4D(Gi).mCherry and MrgprD-mCherry mice were tested before injection. Then the mice were anesthetized with 1% isoflurane and 0.1 mg/ml CNO was injected intraperitoneally at a dose of 1mg/kg body weight. Half an hour after injection, the same behavioral test was repeated.

### Statistics

Data are presented as mean ± SEM, unless otherwise indicated. Multiple groups were compared using One-way or Two-way ANOVA followed by post-hoc tests as indicated in figure legends. Two-tailed *t-tests* were used to compare two groups. The difference was considered to be significant when *p* < 0.05, and ‘#’ stands for ‘not significant’.

## ACKNOWLEDGEMENTS

We thank Genevieve Brindle for the intrathecal injection. This work was supported by Canadian Institutes of Health Research grant FDN 159906 to Y.D.K., Natural Science and Engineering Research Council Discovery grant CG139292 to F. W., and by a Canada Research Chair on Chronic Pain and Related Brain Disorders to Y.D.K. .

## AUTHOR CONTRIBUTIONS

F.W. and Y.D.K. designed the study. F.W. conducted all the imaging experiments. F.W. and S.F. performed all the behavioral experiments. F.W., L-E.L., and P.L. did the immunostaining experiment. CL.L performed the single-cell PCR experiment. E.B. and D.C. designed and built the two-photon microscope and rigged both thermal and mechanical stimulators. S.L.C. designed the Spike2 interface to analyze Ca^2+^ curves. S.L.C. E.B. and A.G. designed and wrote algorithms to parse and analyze the data. B.D. provided the mouse line targeting LTMR and contributed to the imaging experiment. M.E.P. contributed to the viral transduction. F.W., L-E.L., and Y.D.K. analyzed the data. F.W. and Y.D.K. wrote the paper. Y.D.K. supervised the research.

## Reference

1 Raja, S. N. et al. The revised International Association for the Study of Pain definition of pain: concepts, challenges, and compromises. Pain 161, 1976–1982, doi:10.1097/j.pain.0000000000001939 (2020).

2 Woolf, C. J. What is this thing called pain? J Clin Invest 120, 3742–3744, doi:10.1172/JCI45178 (2010).

3 Costigan, M., Scholz, J. & Woolf, C. J. Neuropathic pain: a maladaptive response of the nervous system to damage. Annu Rev Neurosci 32, 1–32, doi:10.1146/annurev.neuro.051508.135531 (2009).

4 Kuner, R. & Flor, H. Structural plasticity and reorganisation in chronic pain. Nat Rev Neurosci 18, 20–30, doi:10.1038/nrn.2016.162 (2016).

5 Treede, R. D. et al. A classification of chronic pain for ICD-11. Pain 156, 1003–1007, doi:10.1097/j.pain.0000000000000160 (2015).

6 Mogil, J. S. Animal models of pain: progress and challenges. Nat Rev Neurosci 10, 283–294, doi:10.1038/nrn2606 (2009).

7 Le Bars, D., Gozariu, M. & Cadden, S. W. Animal models of nociception. Pharmacol Rev 53, 597–652 (2001).

8 Baron, R. et al. Peripheral neuropathic pain: a mechanism-related organizing principle based on sensory profiles. Pain 158, 261–272, doi:10.1097/j.pain.0000000000000753 (2017).

9 Le Pichon, C. E. & Chesler, A. T. The functional and anatomical dissection of somatosensory subpopulations using mouse genetics. Front Neuroanat 8, 21, doi:10.3389/fnana.2014.00021 (2014).

10 Scherrer, G. et al. Dissociation of the opioid receptor mechanisms that control mechanical and heat pain. Cell 137, 1148–1159, doi:10.1016/j.cell.2009.04.019 (2009).

11 Meltzer, S., Santiago, C., Sharma, N. & Ginty, D. D. The cellular and molecular basis of somatosensory neuron development. Neuron 109, 3736–3757, doi:10.1016/j.neuron.2021.09.004 (2021).

12 Zhang, X. & Bao, L. The development and modulation of nociceptive circuitry. Curr Opin Neurobiol 16, 460–466, doi:10.1016/j.conb.2006.06.002 (2006).

13 Cavanaugh, D. J. et al. Distinct subsets of unmyelinated primary sensory fibers mediate behavioral responses to noxious thermal and mechanical stimuli. Proc Natl Acad Sci U S A 106, 9075–9080, doi:10.1073/pnas.0901507106 (2009).

14 Mishra, S. K. & Hoon, M. A. Ablation of TrpV1 neurons reveals their selective role in thermal pain sensation. Mol Cell Neurosci 43, 157–163, doi:10.1016/j.mcn.2009.10.006 (2010).

15 Mishra, S. K., Tisel, S. M., Orestes, P., Bhangoo, S. K. & Hoon, M. A. TRPV1-lineage neurons are required for thermal sensation. EMBO J 30, 582–593, doi:10.1038/emboj.2010.325 (2011).

16 Beaudry, H., Daou, I., Ase, A. R., Ribeiro-da-Silva, A. & Seguela, P. Distinct behavioral responses evoked by selective optogenetic stimulation of the major TRPV1+ and MrgD+ subsets of C-fibers. Pain 158, 2329–2339, doi:10.1097/j.pain.0000000000001016 (2017).

17 Perl, E. R. Ideas about pain, a historical view. Nat Rev Neurosci 8, 71–80, doi:10.1038/nrn2042 (2007).

18 Prescott, S. A., Ma, Q. & De Koninck, Y. Normal and abnormal coding of somatosensory stimuli causing pain. Nat Neurosci 17, 183–191, doi:10.1038/nn.3629 (2014).

19 Ma, Q. Labeled lines meet and talk: population coding of somatic sensations. J Clin Invest 120, 3773–3778, doi:10.1172/JCI43426 (2010).

20 Woodbury, C. J. et al. Nociceptors lacking TRPV1 and TRPV2 have normal heat responses. J Neurosci 24, 6410–6415, doi:10.1523/JNEUROSCI.1421-04.2004 (2004).

21 Lawson, J. J., McIlwrath, S. L., Woodbury, C. J., Davis, B. M. & Koerber, H. R. TRPV1 unlike TRPV2 is restricted to a subset of mechanically insensitive cutaneous nociceptors responding to heat. J Pain 9, 298–308, doi:10.1016/j.jpain.2007.12.001 (2008).

22 Cain, D. M., Khasabov, S. G. & Simone, D. A. Response properties of mechanoreceptors and nociceptors in mouse glabrous skin: an in vivo study. J Neurophysiol 85, 1561–1574, doi:10.1152/jn.2001.85.4.1561 (2001).

23 Koltzenburg, M., Stucky, C. L. & Lewin, G. R. Receptive properties of mouse sensory neurons innervating hairy skin. J Neurophysiol 78, 1841–1850, doi:10.1152/jn.1997.78.4.1841 (1997).

24 Chisholm, K. I., Khovanov, N., Lopes, D. M., La Russa, F. & McMahon, S. B. Large Scale In Vivo Recording of Sensory Neuron Activity with GCaMP6. eNeuro 5, doi:10.1523/ENEURO.0417-17.2018 (2018).

25 Wang, F. et al. Sensory Afferents Use Different Coding Strategies for Heat and Cold. Cell Rep 23, 2001–2013, doi:10.1016/j.celrep.2018.04.065 (2018).

26 Perl, E. R. Cutaneous polymodal receptors: characteristics and plasticity. Prog Brain Res 113, 21–37, doi:10.1016/s0079-6123(08)61079-1 (1996).

27 Prescott, S. A. & Ratte, S. Pain processing by spinal microcircuits: afferent combinatorics. Curr Opin Neurobiol 22, 631–639, doi:10.1016/j.conb.2012.02.010 (2012).

28 Chen, T. W. et al. Ultrasensitive fluorescent proteins for imaging neuronal activity. Nature 499, 295–300, doi:10.1038/nature12354 (2013).

29 Cavanaugh, D. J. et al. Restriction of transient receptor potential vanilloid-1 to the peptidergic subset of primary afferent neurons follows its developmental downregulation in nonpeptidergic neurons. J Neurosci 31, 10119–10127, doi:10.1523/JNEUROSCI.1299-11.2011 (2011).

30 Coste, B. et al. Piezo1 and Piezo2 are essential components of distinct mechanically activated cation channels. Science 330, 55–60, doi:10.1126/science.1193270 (2010).

31 Murthy, S. E. et al. The mechanosensitive ion channel Piezo2 mediates sensitivity to mechanical pain in mice. Sci Transl Med 10, doi:10.1126/scitranslmed.aat9897 (2018).

32 Ranade, S. S. et al. Piezo2 is the major transducer of mechanical forces for touch sensation in mice. Nature 516, 121–125, doi:10.1038/nature13980 (2014).

33 Mogil, J. S. et al. Screening for pain phenotypes: analysis of three congenic mouse strains on a battery of nine nociceptive assays. Pain 126, 24–34, doi:10.1016/j.pain.2006.06.004 (2006).

34 Rau, K. K. et al. Mrgprd enhances excitability in specific populations of cutaneous murine polymodal nociceptors. J Neurosci 29, 8612–8619, doi:10.1523/JNEUROSCI.1057-09.2009 (2009).

35 Dong, X., Han, S., Zylka, M. J., Simon, M. I. & Anderson, D. J. A diverse family of GPCRs expressed in specific subsets of nociceptive sensory neurons. Cell 106, 619–632, doi:10.1016/s0092-8674(01)00483-4 (2001).

36 Zylka, M. J., Rice, F. L. & Anderson, D. J. Topographically distinct epidermal nociceptive circuits revealed by axonal tracers targeted to Mrgprd. Neuron 45, 17–25, doi:10.1016/j.neuron.2004.12.015 (2005).

37 Lee, S. et al. Novel charged sodium and calcium channel inhibitor active against neurogenic inflammation. Elife 8, doi:10.7554/eLife.48118 (2019).

38 Puopolo, M. et al. Permeation and block of TRPV1 channels by the cationic lidocaine derivative QX-314. J Neurophysiol 109, 1704–1712, doi:10.1152/jn.00012.2013 (2013).

39 Roberson, D. P. et al. Activity-dependent silencing reveals functionally distinct itch-generating sensory neurons. Nat Neurosci 16, 910–918, doi:10.1038/nn.3404 (2013).

40 Kim, Y. H. et al. TRPV1 in GABAergic interneurons mediates neuropathic mechanical allodynia and disinhibition of the nociceptive circuitry in the spinal cord. Neuron 74, 640–647, doi:10.1016/j.neuron.2012.02.039 (2012).

41 Pecze, L. et al. Resiniferatoxin mediated ablation of TRPV1+ neurons removes TRPA1 as well. Can J Neurol Sci 36, 234–241, doi:10.1017/s0317167100006600 (2009).

42 Kaufman, D. L., McGinnis, J. F., Krieger, N. R. & Tobin, A. J. Brain glutamate decarboxylase cloned in lambda gt-11: fusion protein produces gamma-aminobutyric acid. Science 232, 1138–1140, doi:10.1126/science.3518061 (1986).

43 Tillakaratne, N. J., Medina-Kauwe, L. & Gibson, K. M. gamma-Aminobutyric acid (GABA) metabolism in mammalian neural and nonneural tissues. Comp Biochem Physiol A Physiol 112, 247–263, doi:10.1016/0300-9629(95)00099-2 (1995).

44 Lorenzo, L. E. et al. Spatial and temporal pattern of changes in the number of GAD65-immunoreactive inhibitory terminals in the rat superficial dorsal horn following peripheral nerve injury. Mol Pain 10, 57, doi:10.1186/1744-8069-10-57 (2014).

45 Lorenzo, L. E. et al. Enhancing neuronal chloride extrusion rescues alpha2/alpha3 GABA(A)-mediated analgesia in neuropathic pain. Nat Commun 11, 869, doi:10.1038/s41467-019-14154-6 (2020).

46 Rivera, C. et al. The K+/Cl-co-transporter KCC2 renders GABA hyperpolarizing during neuronal maturation. Nature 397, 251–255, doi:10.1038/16697 (1999).

47 Virtanen, M. A., Uvarov, P., Mavrovic, M., Poncer, J. C. & Kaila, K. The Multifaceted Roles of KCC2 in Cortical Development. Trends Neurosci 44, 378–392, doi:10.1016/j.tins.2021.01.004 (2021).

48 Delpire, E. Cation-Chloride Cotransporters in Neuronal Communication. News Physiol Sci 15, 309–312, doi:10.1152/physiologyonline.2000.15.6.309 (2000).

49 Doyon, N., Vinay, L., Prescott, S. A. & De Koninck, Y. Chloride Regulation: A Dynamic Equilibrium Crucial for Synaptic Inhibition. Neuron 89, 1157–1172, doi:10.1016/j.neuron.2016.02.030 (2016).

50 Chamma, I., Chevy, Q., Poncer, J. C. & Levi, S. Role of the neuronal K-Cl co-transporter KCC2 in inhibitory and excitatory neurotransmission. Front Cell Neurosci 6, 5, doi:10.3389/fncel.2012.00005 (2012).

51 Kahle, K. T. et al. Modulation of neuronal activity by phosphorylation of the K-Cl cotransporter KCC2. Trends Neurosci 36, 726–737, doi:10.1016/j.tins.2013.08.006 (2013).

52 Ferrini, F. et al. Differential chloride homeostasis in the spinal dorsal horn locally shapes synaptic metaplasticity and modality-specific sensitization. Nat Commun 11, 3935, doi:10.1038/s41467-020-17824-y (2020).

53 Sternson, S. M. & Roth, B. L. Chemogenetic tools to interrogate brain functions. Annu Rev Neurosci 37, 387–407, doi:10.1146/annurev-neuro-071013-014048 (2014).

54 Wang, H. & Zylka, M. J. Mrgprd-expressing polymodal nociceptive neurons innervate most known classes of substantia gelatinosa neurons. J Neurosci 29, 13202–13209, doi:10.1523/JNEUROSCI.3248-09.2009 (2009).

55 Warwick, C. et al. MrgprdCre lineage neurons mediate optogenetic allodynia through an emergent polysynaptic circuit. Pain 162, 2120–2131, doi:10.1097/j.pain.0000000000002227 (2021).

56 Qi, L. et al. A mouse DRG genetic toolkit reveals morphological and physiological diversity of somatosensory neuron subtypes. Cell 187, 1508–1526 e1516, doi:10.1016/j.cell.2024.02.006 (2024).

57 Wang, H. et al. A novel spinal neuron connection for heat sensation. Neuron 110, 2315–2333 e2316, doi:10.1016/j.neuron.2022.04.021 (2022).

58 Braz, J., Solorzano, C., Wang, X. & Basbaum, A. I. Transmitting pain and itch messages: a contemporary view of the spinal cord circuits that generate gate control. Neuron 82, 522–536, doi:10.1016/j.neuron.2014.01.018 (2014).

59 Warwick, C. et al. Cell type-specific calcium imaging of central sensitization in mouse dorsal horn. Nat Commun 13, 5199, doi:10.1038/s41467-022-32608-2 (2022).

60 Ganley, R. P. et al. Inhibitory Interneurons That Express GFP in the PrP-GFP Mouse Spinal Cord Are Morphologically Heterogeneous, Innervated by Several Classes of Primary Afferent and Include Lamina I Projection Neurons among Their Postsynaptic Targets. J Neurosci 35, 7626–7642, doi:10.1523/JNEUROSCI.0406-15.2015 (2015).

61 Craig, A. D., Krout, K. & Andrew, D. Quantitative response characteristics of thermoreceptive and nociceptive lamina I spinothalamic neurons in the cat. J Neurophysiol 86, 1459–1480, doi:10.1152/jn.2001.86.3.1459 (2001).

62 Polgar, E., Al Ghamdi, K. S. & Todd, A. J. Two populations of neurokinin 1 receptor-expressing projection neurons in lamina I of the rat spinal cord that differ in AMPA receptor subunit composition and density of excitatory synaptic input. Neuroscience 167, 1192–1204, doi:10.1016/j.neuroscience.2010.03.028 (2010).

63 Cordero-Erausquin, M. et al. Dorsal horn neurons presynaptic to lamina I spinoparabrachial neurons revealed by transynaptic labeling. J Comp Neurol 517, 601–615, doi:10.1002/cne.22179 (2009).

64 Lu, Y. & Perl, E. R. Modular organization of excitatory circuits between neurons of the spinal superficial dorsal horn (laminae I and II). J Neurosci 25, 3900–3907, doi:10.1523/JNEUROSCI.0102-05.2005 (2005).

65 Wang, L. B. et al. Parallel Spinal Pathways for Transmitting Reflexive and Affective Dimensions of Nocifensive Behaviors Evoked by Selective Activation of the Mas-Related G Protein-Coupled Receptor D-Positive and Transient Receptor Potential Vanilloid 1-Positive Subsets of Nociceptors. Front Cell Neurosci 16, 910670, doi:10.3389/fncel.2022.910670 (2022).

66 Duan, B. et al. Identification of spinal circuits transmitting and gating mechanical pain. Cell 159, 1417–1432, doi:10.1016/j.cell.2014.11.003 (2014).

67 Christensen, A. J. et al. In Vivo Interrogation of Spinal Mechanosensory Circuits. Cell Rep 17, 1699–1710, doi:10.1016/j.celrep.2016.10.010 (2016).

68 Li, B. et al. A novel analgesic approach to optogenetically and specifically inhibit pain transmission using TRPV1 promoter. Brain Res 1609, 12–20, doi:10.1016/j.brainres.2015.03.008 (2015).

69 Ma, Q. A functional subdivision within the somatosensory system and its implications for pain research. Neuron 110, 749–769, doi:10.1016/j.neuron.2021.12.015 (2022).

70 Huang, T. et al. Identifying the pathways required for coping behaviours associated with sustained pain. Nature 565, 86–90, doi:10.1038/s41586-018-0793-8 (2019).

71 Magnusson, B. M. & Koskinen, L. D. In vitro percutaneous penetration of topically applied capsaicin in relation to in vivo sensation responses. Int J Pharm 195, 55–62, doi:10.1016/s0378-5173(99)00337-3 (2000).

72 Klein, A. et al. Pruriception and neuronal coding in nociceptor subtypes in human and nonhuman primates. Elife 10, doi:10.7554/eLife.64506 (2021).

73 Li, L. et al. The functional organization of cutaneous low-threshold mechanosensory neurons. Cell 147, 1615–1627, doi:10.1016/j.cell.2011.11.027 (2011).

74 Desiderio, S. et al. Touch receptor end-organ innervation and function require sensory neuron expression of the transcription factor Meis2. Elife 12, doi:10.7554/eLife.89287 (2024).

75 Guo, A., Vulchanova, L., Wang, J., Li, X. & Elde, R. Immunocytochemical localization of the vanilloid receptor 1 (VR1): relationship to neuropeptides, the P2X3 purinoceptor and IB4 binding sites. Eur J Neurosci 11, 946–958, doi:10.1046/j.1460-9568.1999.00503.x (1999).

76 Tominaga, M. et al. The cloned capsaicin receptor integrates multiple pain-producing stimuli. Neuron 21, 531–543, doi:10.1016/s0896-6273(00)80564-4 (1998).

77 Ribeiro-da-Silva, A. & De Koninck, Y. in The Senses: A Comprehensive Reference (eds Richard H. Masland et al.) 279-310 (Academic Press, 2008).

78 Papalampropoulou-Tsiridou, M. et al. Distribution of acid-sensing ion channel subunits in human sensory neurons contrasts with that in rodents. Brain Commun 4, fcac256, doi:10.1093/braincomms/fcac256 (2022).

79 Shiers, S., Klein, R. M. & Price, T. J. Quantitative differences in neuronal subpopulations between mouse and human dorsal root ganglia demonstrated with RNAscope in situ hybridization. Pain 161, 2410–2424, doi:10.1097/j.pain.0000000000001973 (2020).

80 Kumazawa, T. & Perl, E. R. Primate cutaneous sensory units with unmyelinated (C) afferent fibers. J Neurophysiol 40, 1325–1338, doi:10.1152/jn.1977.40.6.1325 (1977).

81 Torebjork, H. E. & Ochoa, J. L. New method to identify nociceptor units innervating glabrous skin of the human hand. Exp Brain Res 81, 509–514, doi:10.1007/BF02423499 (1990).

82 Torebjork, H. E. Afferent C units responding to mechanical, thermal and chemical stimuli in human non-glabrous skin. Acta Physiol Scand 92, 374–390, doi:10.1111/j.1748-1716.1974.tb05755.x (1974).

83 Lorenzo, L. E. et al. Gephyrin clusters are absent from small diameter primary afferent terminals despite the presence of GABA(A) receptors. J Neurosci 34, 8300–8317, doi:10.1523/JNEUROSCI.0159-14.2014 (2014).

84 Li, C. L. et al. Somatosensory neuron types identified by high-coverage single-cell RNA-sequencing and functional heterogeneity. Cell Res 26, 967, doi:10.1038/cr.2016.90 (2016).

85 Davalos, D. et al. Stable in vivo imaging of densely populated glia, axons and blood vessels in the mouse spinal cord using two-photon microscopy. J Neurosci Methods 169, 1–7, doi:10.1016/j.jneumeth.2007.11.011 (2008).

86 Vrontou, S., Wong, A. M., Rau, K. K., Koerber, H. R. & Anderson, D. J. Genetic identification of C fibres that detect massage-like stroking of hairy skin in vivo. Nature 493, 669–673, doi:10.1038/nature11810 (2013).

87 Veilleux, I., Spencer, J. A., Biss, D. P., Cote, D. & Lin, C. P. J. I. J. o. s. t. i. q. e. In vivo cell tracking with video rate multimodality laser scanning microscopy. 14, 10–18 (2008).

