## Supplemental Table and Figure legend for "Thermal and mechanical modalities converge in the noxious range"

**Table 1 Primary antibodies used in this study**

| Antibody | Species | Source | Cat.No. | Dilution |
|---|---|---|---|---|
| TRPV1 | Goat | Santa Cruz | sc-12498 | 1:1,000 |
| TRPV1 | Rabbit | Alomone Labs | ACC-030 | 1:6,000 |
| CGRP | Rabbit | Sigma Aldrich | C8198 | 1:2,000 |
| CGRP | Mouse | Sigma Aldrich | C7113 | 1:2,000 |
| GFP | Mouse | Invitrogen | A-11120 | 1:500 |
| NF200 | Chicken | Abcam | Ab4680 | 1:2,000 |
| P2X3 | Rabbit | Neuromics | RA10109 | 1:2,000 |
| GAD65/67 | Rabbit | Sigma Aldrich | G5163 | 1:1,000 |
| KCC2 | Rabbit | Sigma Aldrich | 07-432 | 1:1,000 |
| NeuN | Chicken | Sigma Aldrich | ABN91 | 1:1,000 |

**Table 2 Secondary antibodies and dyes used in this study**

| Antibody/dye | Source | Cat.No. | Dilution |
|---|---|---|---|
| Donkey-anti-Goat, Cy5 | Jackson Immuno Research | 705-175-147 | 1:200 |
| Donkey-anti-Rabbit, Rhodamine Red | Jackson Immuno Research | 711-295-152 | 1:200 |
| Goat-anti-Rabbit, Cy3 | Jackson Immuno Research | 111-165-144 | 1:500 |
| Goat-anti-Mouse Alexa Fluor 488 | Invitrogen | A-11001 | 1:500 |
| Goat-anti-Chicken Alexa Fluor 647 | Invitrogen | A-21449 | 1:500 |
| Goat-anti-Chicken Alexa Fluor 546 | Invitrogen | A-11040 | 1:200 |
| Goat-anti-Rabbit Alexa Fluor 647 | Invitrogen | A-21245 | 1:500 |
| IB4 conjugated to Alexa Fluor 488 | Invitrogen | I21411 | 1:250 |
| IB4 conjugated to Alexa Fluor 568 | Invitrogen | I21412 | 1:250 |
| IB4 conjugated to Alexa Fluor 647 | Invitrogen | I32450 | 1:250 |

### Supplemental FIGURE LEGENDS

#### Figure S1. Validation of Cre-dependent GCaMP6s virus

Representative micrographs of DRG sections showing Cre-dependent GCaMP6s virus were only expressed in the DRGs of $Cre^+$ mice (left), but not the wild-type mice (right). Scale bar = 100 μm.

#### Figure S2. TRPV1 is expressed in the majority of peptidergic neurons in adult mice

(A) Immunostaining results showing the colocalization of $TRPV1^+$, $CGRP^+$, and $IB4^+$ DRG neurons in adult wild-type mice. Yellow arrows pointed to TRPV1/CGRP double-positive (IB4-negative) neurons. Scale bar = 50 μm.

(B) Euler diagrams showed that the majority of $TRPV1^+$ neurons expressed CGRP, but didn't bind to IB4.

(C) Immunostaining results showing the colocalization of $TRPV1^+$, $P2X3^+$, and $IB4^+$ DRG neurons in adult wild-type mice. Yellow arrows pointed to TRPV1/P2X3 double-positive (IB4-negative) neurons.

(D) Euler diagrams showed that the majority of $TRPV1^+$ neurons did not express P2X3.

#### Figure S3. Expression of Cre-dependent GCaMP6s virus in $TRPV1^{Cre/Cre}$ mice

(A) Representative micrographs of DRG sections showing GCaMP6s expression (without antibody amplification) superimposed on labeling of TRPV1 and IB4 in $TRPV1^{Cre/Cre}$ mice. Cyan arrows pointed to GCaMP/IB4 double-positive (TRPV1-negative) neurons. Scale bar = 50 μm.

(B) Euler diagrams showed that a substantial subpopulation (28%) of $GCaMP^+$ neurons do not express TRPV1 and they are non-peptidergic in $TRPV1^{Cre/Cre}$ mice.

#### Figure S4. Characterization of pinch-sensitive and pinch-insensitive $TRPV1^+$ thermoreceptors

(A) The soma size of pinch-sensitive and pinch-insensitive $TRPV1^+$ neurons were not different (n = 18 for pinch-insensitive neurons and n = 40 for pinch-sensitive neurons; Welch's unpaired *t* test).

(B-D) While $Ca^{2+}$ responses to 50°C stimulation lasted longer in pinch-sensitive neurons than pinch-insensitive $TRPV1^+$ neurons (C; n = 68 for pinch-insensitive neurons and n = 75 for pinch-sensitive neurons, $p = 0.016$), the peak amplitude (B; n = 68 for pinch-insensitive neurons and n = 75 for pinch-sensitive neurons $p = 0.079$) and decay time (85% to 15% of peak; D; n = 66 for pinch-insensitive neurons and n = 68 for pinch-sensitive neurons, $p = 0.408$) were not different between the two groups by using unpaired *t* test with Welch's correction.

**Figure S5. Expression of Cre-dependent GCaMP6s virus in MrpgrD$^{Cre/+}$ mice**

(A) Representative micrographs of DRG sections showing GCaMP6s expression (without antibody amplification) superimposed on labeling of CGRP in MrgprD$^{Cre/+}$ mice. The yellow arrows pointed to the neurons that were GCaMP6s and CGRP-double positive, while the green and red arrows pointed to the neurons that were only GCaMP6s or CGRP-positive, respectively. Scale bar = 50 µm.

(B) Euler diagrams showed that the majority of GCaMP$^+$ neurons did not express CGRP.

**Figure S6. Decrease of GAD65/67, but not KCC2, in the superficial spinal dorsal horn after i.t. capsaicin-mediated ablation of TRPV1$^+$ afferents**

(A) Representative confocal images of the spinal cord dorsal horn in male mice with intrathecal injection of 10 µg capsaicin or vehicle showing staining of GAD65/67, CGRP, IB4, and NeuN. Scale bar = 33.5 µm.

(B) Representative confocal images of the spinal cord dorsal horn in male mice with intrathecal injection of 10 µg capsaicin or vehicle showing staining of KCC2, CGRP, IB4, and NeuN. Scale bar = 33.5 µm.

(C) Average fluorescence intensity of KCC2 on the neuronal membrane in the dorsal horn as a function of the position towards the IB4 barycentric origin in vehicle i.t. injected (n = 8) vs. capsaicin i.t. injected mice (n = 8).

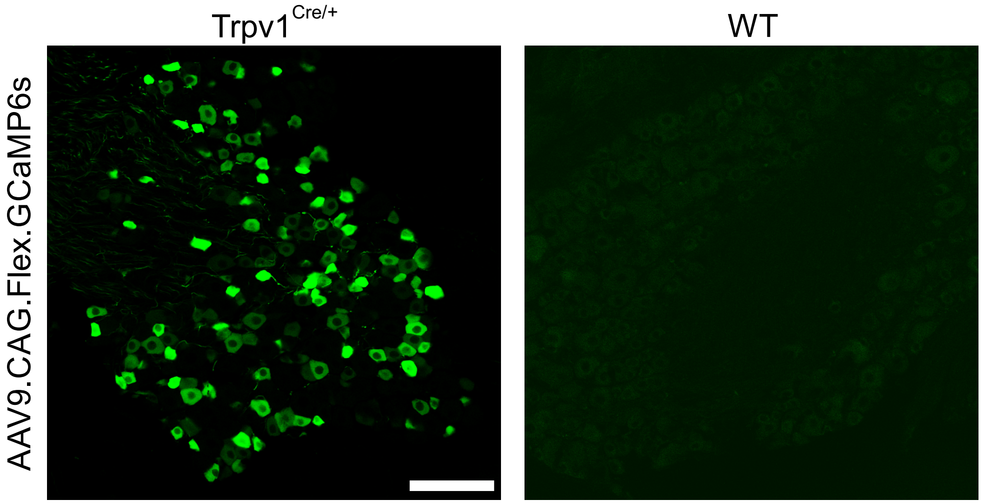
Trpv1^Cre/+
WT
AAV9.CAG.Flex.GCaMP6s

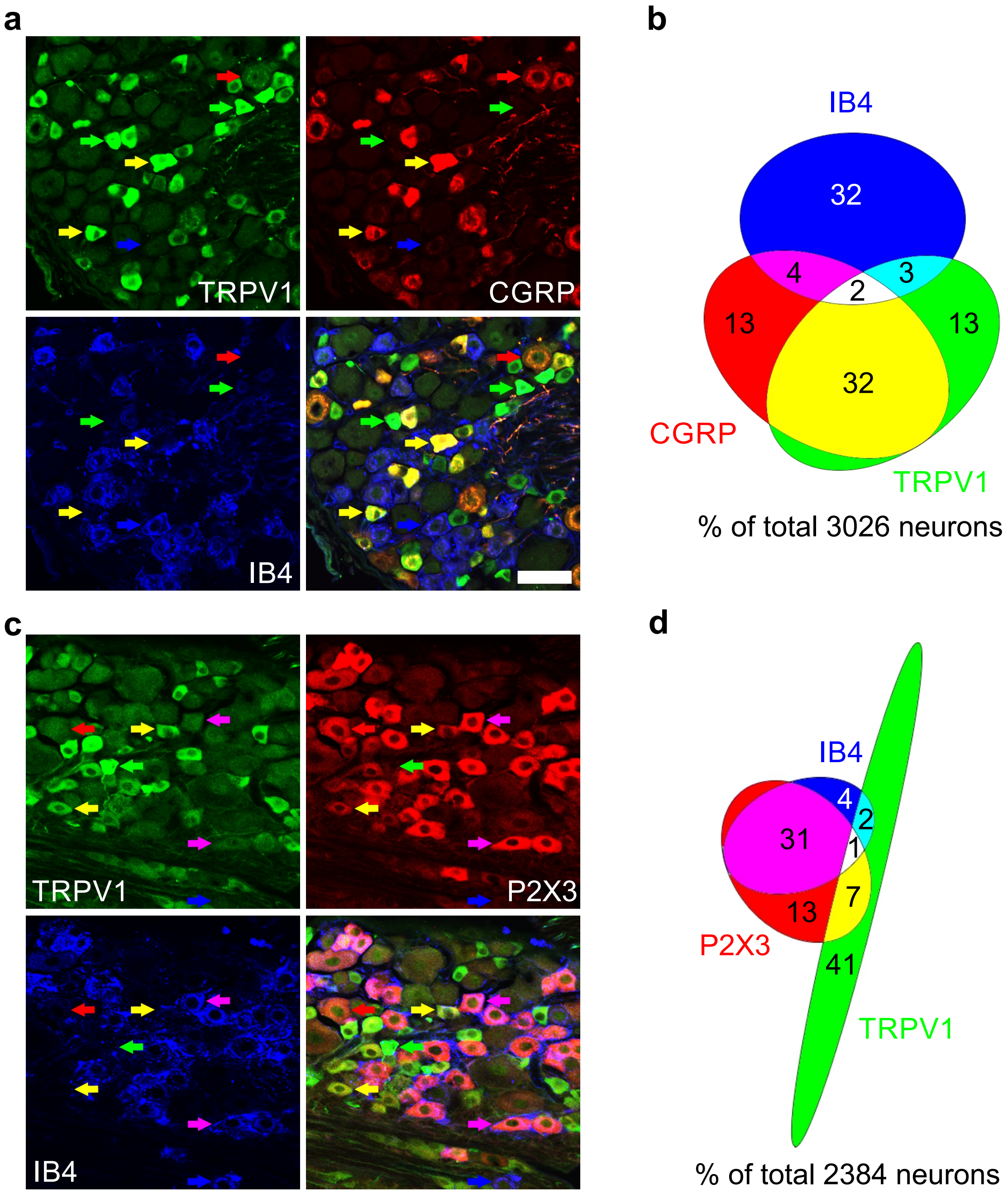

a
TRPV1
CGRP
IB4
b
IB4
32
4
3
2
13
13
32
CGRP
TRPV1
% of total 3026 neurons
c
TRPV1
P2X3
IB4
d
IB4
4
2
31
1
7
13
P2X3
41
TRPV1
% of total 2384 neurons

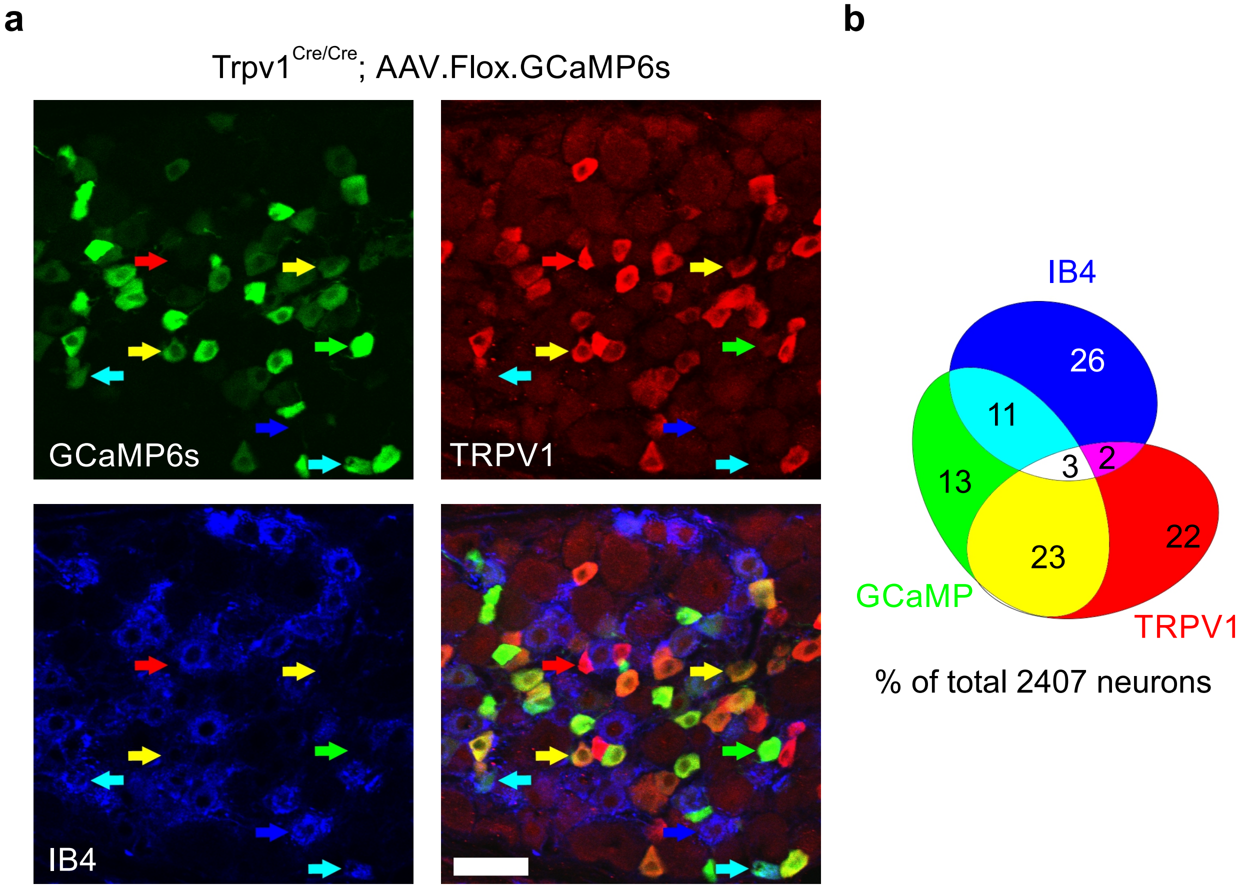
a
Trpv1 Cre/Cre; AAV.Flox.GCaMP6s
GCaMP6s
TRPV1
IB4
b
IB4
26
11
13
3
2
22
23
GCaMP
TRPV1
% of total 2407 neurons

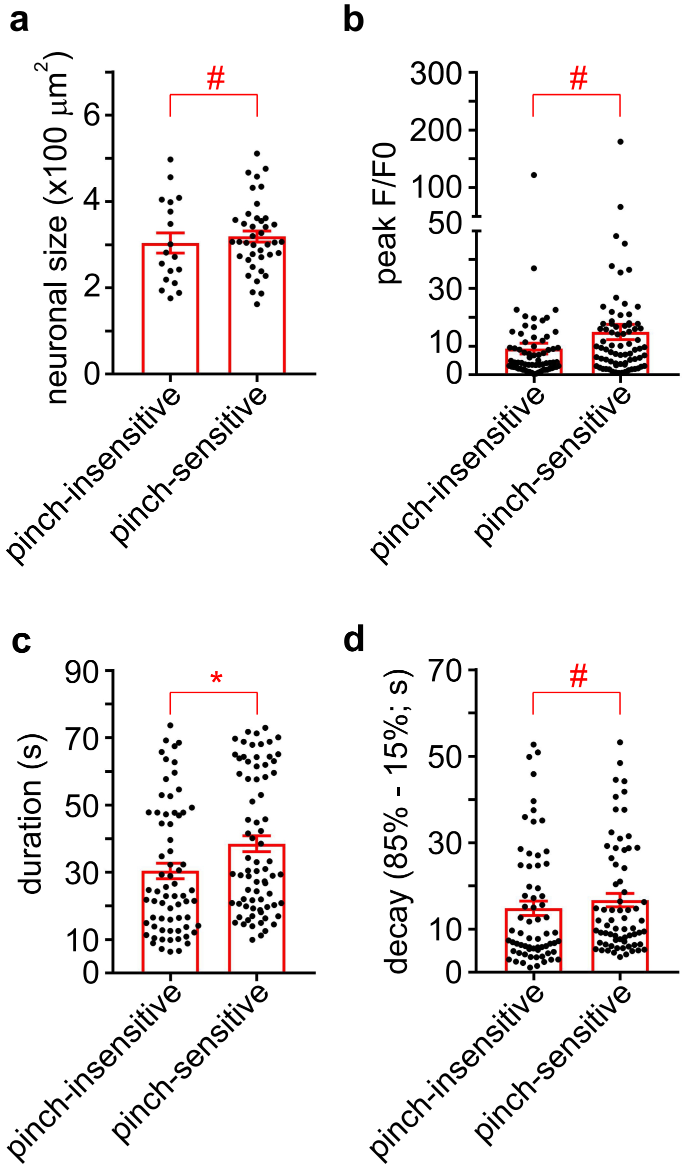

a
b
c
d
neuronal size (x100 μm$^2$)
peak F/F0
duration (s)
decay (85% - 15%; s)
#
*
pinch-insensitive
pinch-sensitive

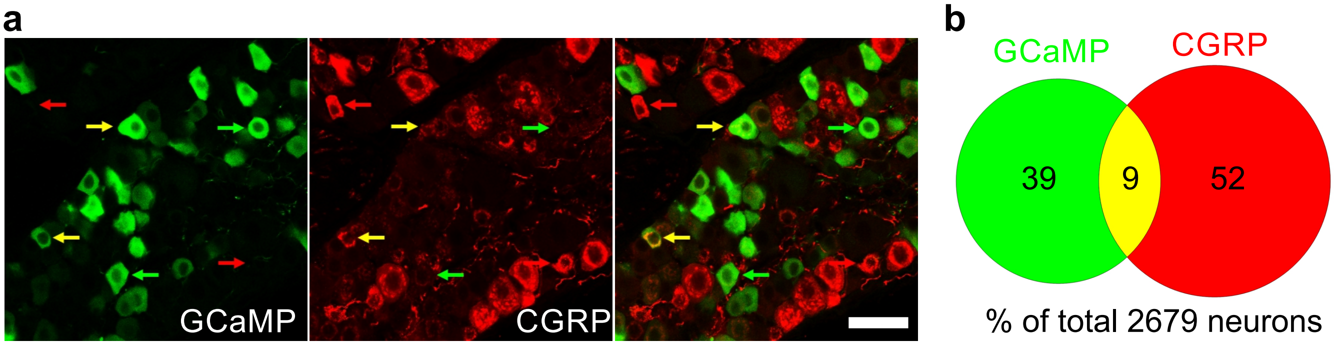
a
GCaMP
CGRP
b
GCaMP
CGRP
39
9
52
% of total 2679 neurons

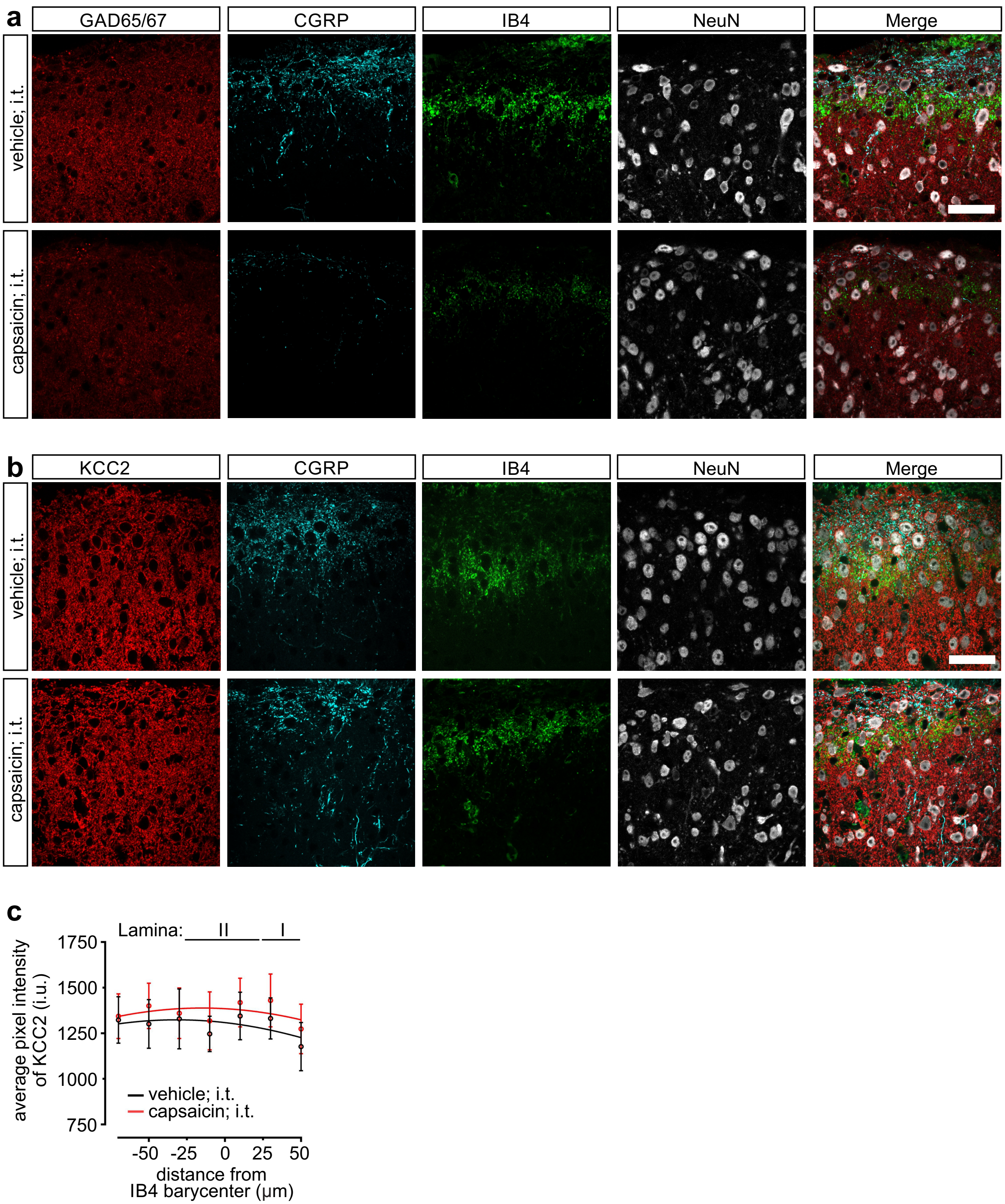
a
GAD65/67
CGRP
IB4
NeuN
Merge
vehicle; i.t.
capsaicin; i.t.
b
KCC2
CGRP
IB4
NeuN
Merge
vehicle; i.t.
capsaicin; i.t.
c
Lamina: II I
average pixel intensity of KCC2 (i.u.)
1750
1500
1250
1000
750
vehicle; i.t.
capsaicin; i.t.
-50 -25 0 25 50
distance from IB4 barycenter (μm)
