## Supplementary figures and images for "Thermal and mechanical modalities converge in the noxious range"

### Supplemental Figures

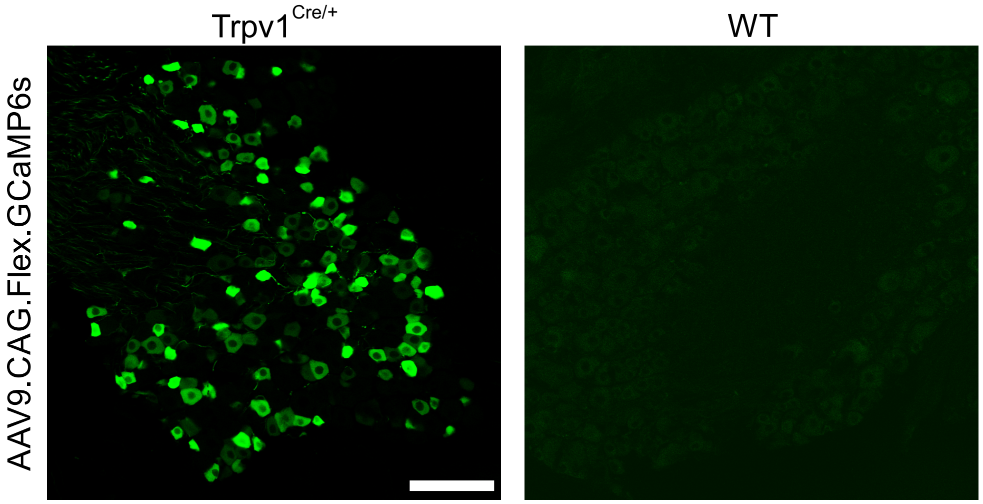

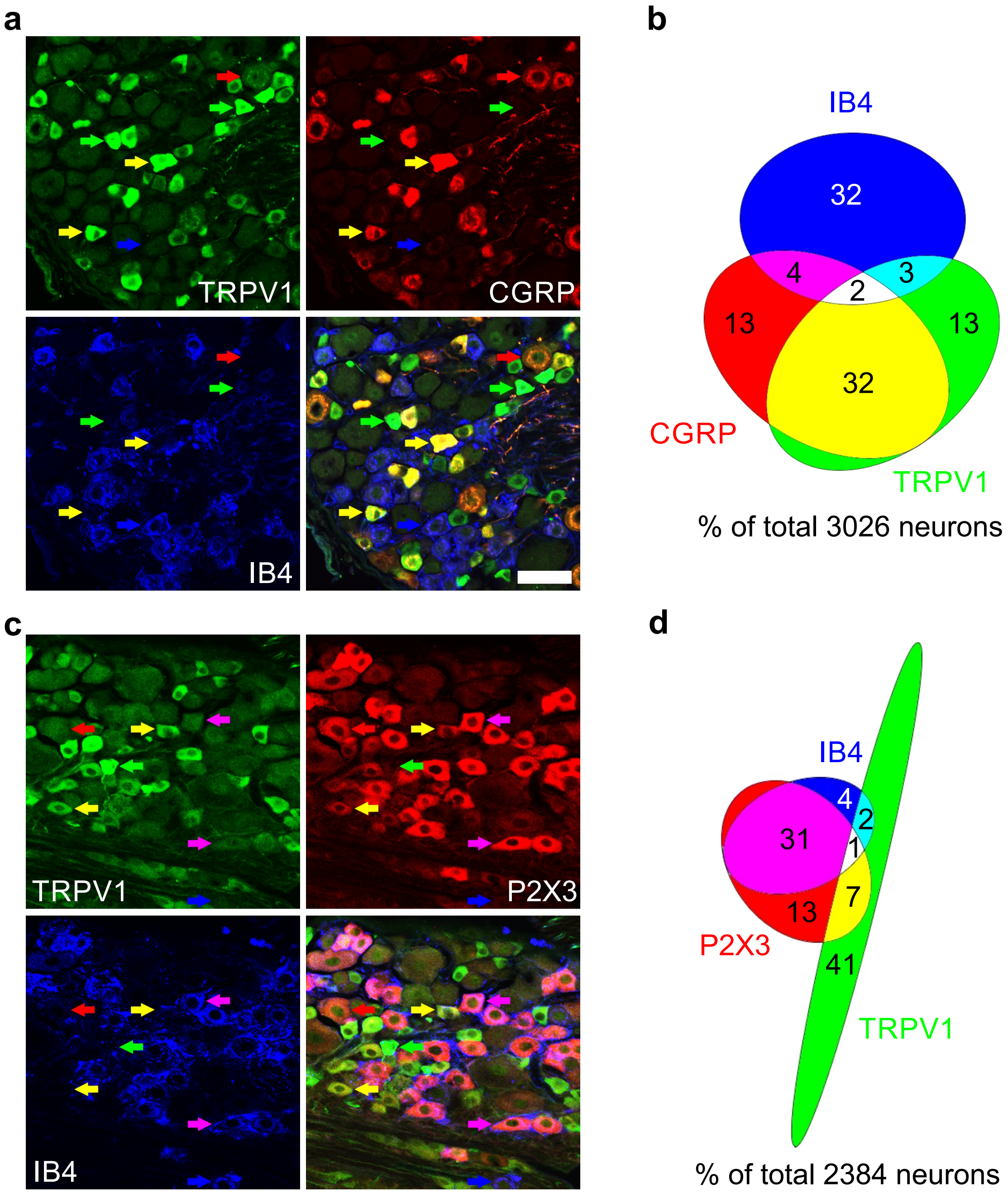

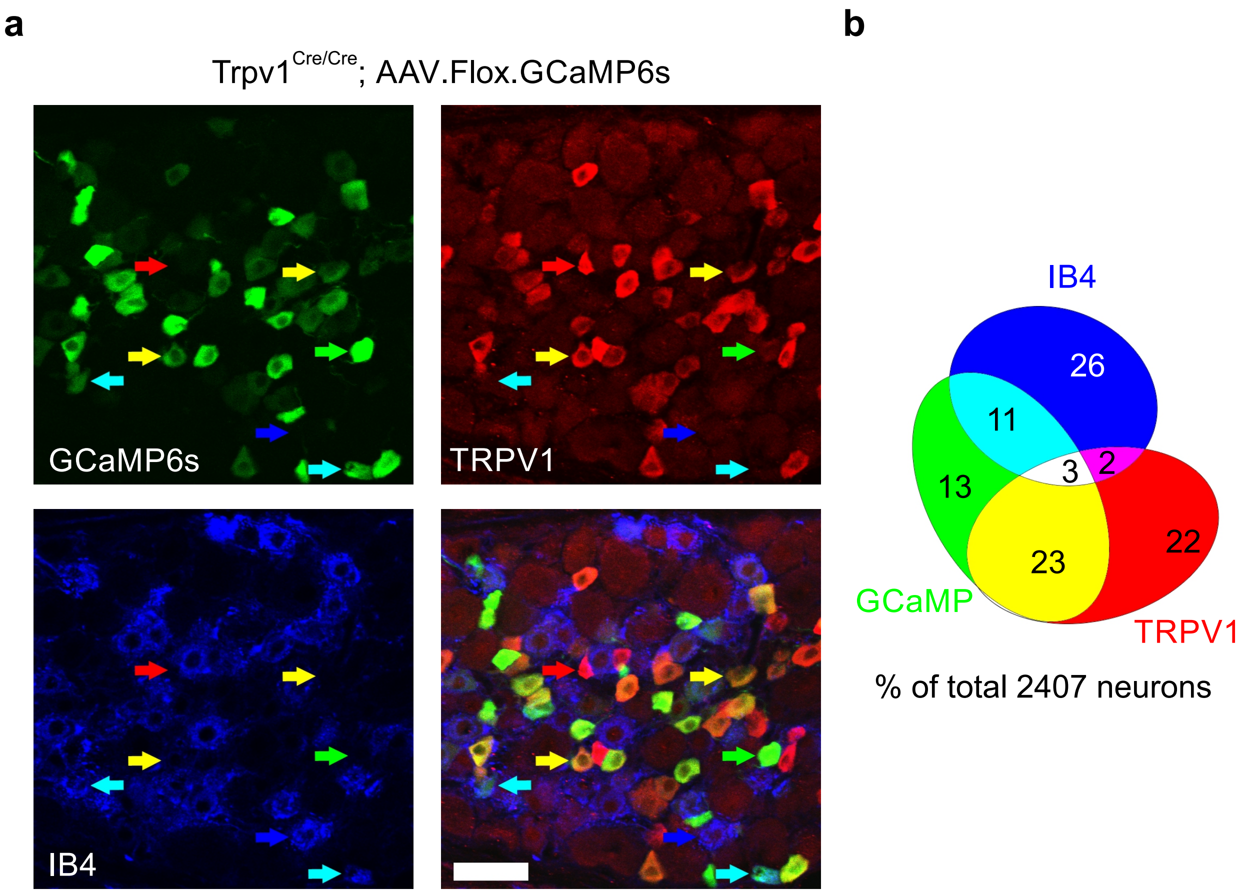

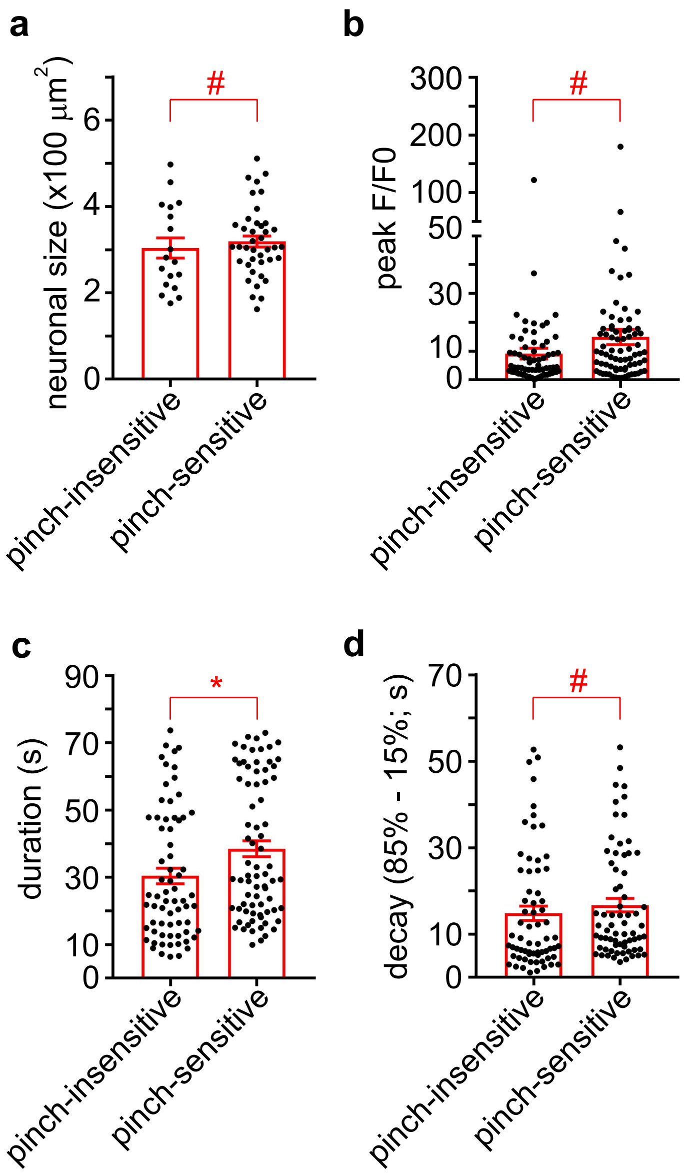

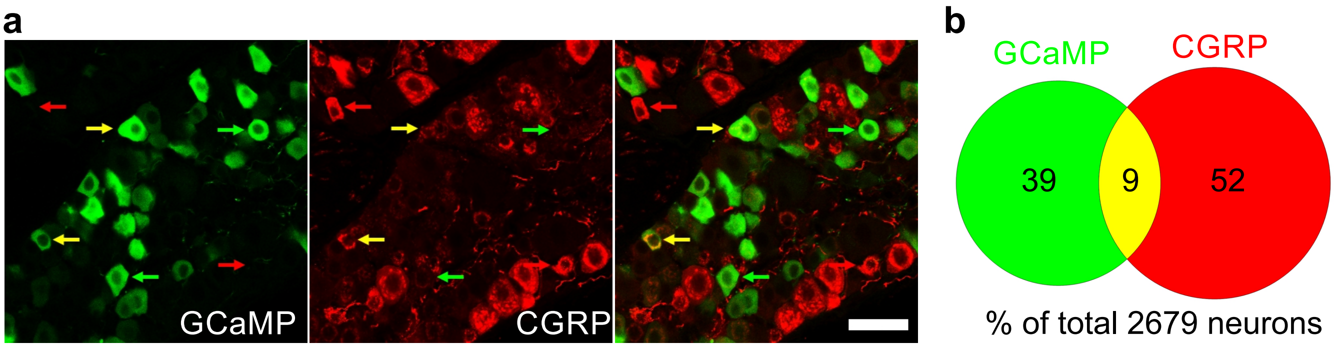

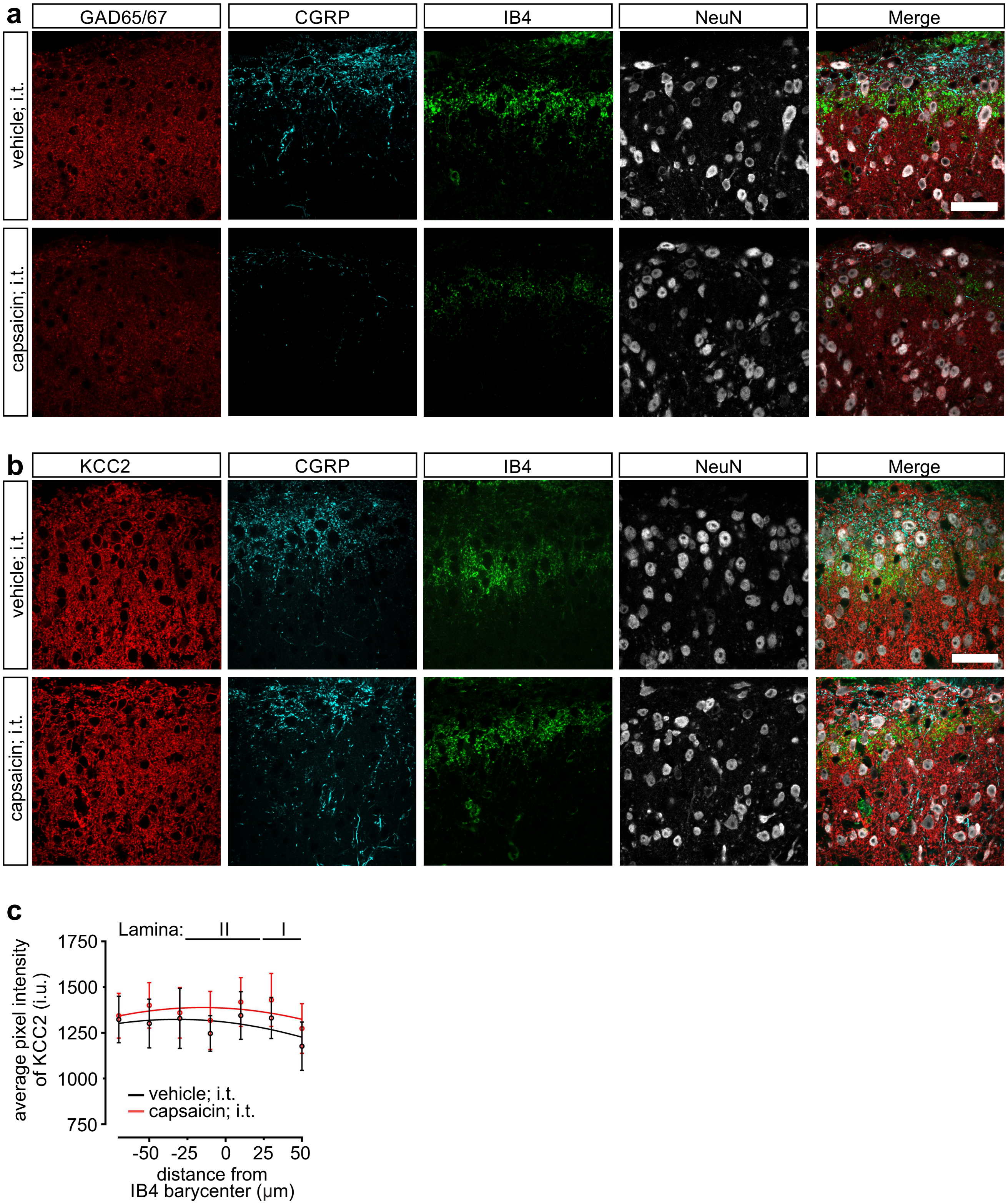
